# Initiation of DNA replication requires actin dynamics and formin activity

**DOI:** 10.1101/102806

**Authors:** Parisis Nikolaos, Liliana Krasinska, Bethany Harker, Serge Urbach, Michel Rossignol, Alain Camasses, James Dewar, Nathalie Morin, Daniel Fisher

**Affiliations:** Montpellier Institute of Molecular Genetics (IGMM), CNRS UMR 5535, 34293 Montpellier, France.; University of Montpellier, Faculty of Sciences, 34090 Montpellier, France.; Present address: Wellcome Trust Centre for Cell Biology, University of Edinburgh, EH9 3QR Edinburgh, UK.; Functional Proteomics Platform (FPP), Institute of Functional Genomics (IGF), CNRS UMR 5203, INSERM U661, 34090 Montpellier, France.; Laboratory of Functional Proteomics, INRA, 34000 Montpellier, France.; Vanderbilt University, Tennessee 37235, USA; Centre de Recherche de Biochimie Macromoléculaire, CNRS UMR5237, 34293 Montpellier, France.; Present address: Institut Jacques Monod, CNRS UMR7592, University Paris Diderot, 75013 Paris, France; Co-first author.

**Keywords:** DNA replication, nuclear transport, actin, formin, CDK

## Abstract

Nuclear actin influences transcription in a manner dependent on its dynamics of polymerisation and nucleocytoplasmic translocation. Using human somatic cells and transcriptionally-silent *Xenopus* egg extracts, we show that actin dynamics is also required for DNA replication. We identify many actin regulators in replicating nuclei from *Xenopus* egg extracts, and show that in human cells, nuclear actin filaments form in early G1 and disassemble prior to S-phase. In either system, treatments that stabilise nuclear actin filaments abrogate nuclear transport and initiation of DNA replication. Mechanistically, actin directly binds RanGTP-importin complexes and disruption of its dynamics hinders cargo release. This prevents both nuclear pore complex (NPC) formation and active nuclear transport, which we show is required throughout DNA replication. Nuclear formin activity is required for two further steps: loading of cyclin-dependent kinase (CDK) and proliferating cell nuclear antigen (PCNA) onto chromatin and initiation of DNA replication. Thus, actin dynamics and formins are involved in several nuclear processes essential for cell proliferation.

## Introduction

In mammalian cells, various functions have been attributed to nuclear actin (Huet *et al*, 2012). Monomeric actin binds chromatin remodeling and RNA polymerase complexes (Rando *et al*, 2002; Kapoor *et al*, 2013) and promotes transcription by all three RNA polymerases (Hofmann *et al*, 2004; Hu *et al*, 2004; Philimonenko *et al*, 2004). Its nuclear levels are regulated by active transport between the nucleus and cytoplasm (Stüven *et al*, 2003; Dopie *et al*, 2012) and polymerisation (Baarlink *et al*, 2013; Lundquist *et al*, 2014; Vartiainen *et al*, 2007). This dynamics is complex: monomeric actin promotes export of the serum response factor (SRF) cofactor MAL/MRTF, extinguishing SRF, yet both nuclear actin polymerisation (Baarlink *et al*, 2013) and depolymerisation (Lundquist *et al*, 2014) can induce SRF-dependent transcription. In epithelial cells, loss of nuclear actin triggers quiescence by disrupting binding of RNA polymerases to their transcription sites (Spencer *et al*, 2011). Nuclear actin also affects co-repressor eviction from promoters (Huang *et al*, 2011) and it can bind to gene regulatory regions (Miyamoto *et al*, 2011; Miyamoto *et al*, 2013a).

*In vitro*, purified actin and profilin self-assemble into long filaments, but cells additionally require actin nucleation factors. Sub-populations of nuclear actin have distinct mobilities, suggesting existence of polymeric forms (Dopie *et al*, 2012; McDonald *et al*, 2006), and several regulators of actin polymerisation have been found in nuclei (Khoudoli *et al*, 2008; Miyamoto *et al*, 2013b; Obrdlik & Percipalle, 2011; Wu *et al*, 2006; Yoo *et al*, 2007; Dopie *et al*, 2015). Physiological nuclear actin polymerisation remains poorly characterised due to difficulties in staining nuclear actin with phalloidin (Grosse & Vartiainen, 2013). In specific settings, like the giant non-replicating nuclei of amphibian oocytes, a filamentous actin network has scaffolding functions (Clark & Rosenbaum, 1979; Gounon & Karsenti, 1981; Feric & Brangwynne, 2013). Stabilised nuclear actin filaments are observed in several pathologies (Lanerolle, 2012) and can be induced by various manipulations, including heat shock and DMSO treatment (Iida *et al*, 1986; Sanger *et al*, 1980); increasing nuclear actin concentrations (Stüven *et al*, 2003; Kalendová *et al*, 2014); activation of nuclear mDia formin (Baarlink *et al*, 2013); overexpression of NLS-tagged IQGAP1 (Johnson *et al*, 2013) or supervillin (Serebryannyy *et al*, 2016); or knockdown of MICAL-2, which promotes nuclear actin depolymerisation through methionine oxidation (Lundquist *et al*, 2014). Stabilisation of nuclear actin filaments inhibits transcription by RNA polymerase II (Serebryannyy *et al*, 2016), whereas serum stimulation of mouse fibroblasts triggers transient nuclear actin filament formation, promoting SRF-dependent transcription (Baarlink *et al*, 2013).

We investigated whether nuclear actin has transcription-independent roles in cell proliferation by using transcriptionally silent *Xenopus* egg extracts (XEE). This system recapitulates early embryonic cell cycles *in vitro*, allowing identification of nuclear assembly pathways (Hetzer *et al*, 2005) and DNA replication mechanisms (Arias & Walter, 2004). Using XEE as well as human somatic cells, we show that actin dynamics is required for initiation of DNA replication, through at least two mechanisms: first, actin dynamics is required for nuclear transport, as polymeric nuclear actin locks cargo-importin-Ran complexes, preventing cargo release from importins. Second, nuclear formin activity promotes chromatin loading of DNA replication factors allowing initiation of DNA synthesis.

## Results

### Nuclear actin dynamics during the cell cycle

We first analysed the combined nucleoskeleton and chromatin proteome of nuclei assembled in XEE by label-free high-resolution mass spectrometry. To assess possible cell cycle regulation of nuclear assembly, we compared replicating nuclei with nuclei assembled in the presence of purvalanol A (PA) to inhibit CDKs (Echalier *et al*, 2012) (Fig 1A). We identified 2610 non-redundant proteins (Fig 1B, C; Supplementary Figure 1, S1). Enriched biological processes included DNA metabolism, chromatin organisation, and, interestingly, regulation of actin polymerisation (Fig 1D; Table S2). We identified 55 actin regulators (Tables S3 & S4), including actin filament nucleating factors such as formins and the Arp2/3 complex. These were unaffected by CDK activity, unlike chromatin recruitment of proteins involved in DNA replication, DNA repair and the S-phase checkpoint (Fig 1B–E; and Tables S1&2). Immunofluorescence analysis confirmed that many actin polymerisation regulators localised to replicating nuclei (Fig 2A), where actin was mostly insoluble (Fig 2B). To visualise nuclear actin directly, we added trace concentrations of fluorescently-labeled actin protein to XEE. This revealed filaments in the egg cytosol, as expected, and both diffuse and patterned intra-nuclear staining (Fig 2C–E). The latter might be actin polymers or monomeric actin associated with other structures, for example chromatin. Labelled DNase1, a high affinity G-actin probe, mainly stained chromatin (Fig 2F).

**Figure 1.**
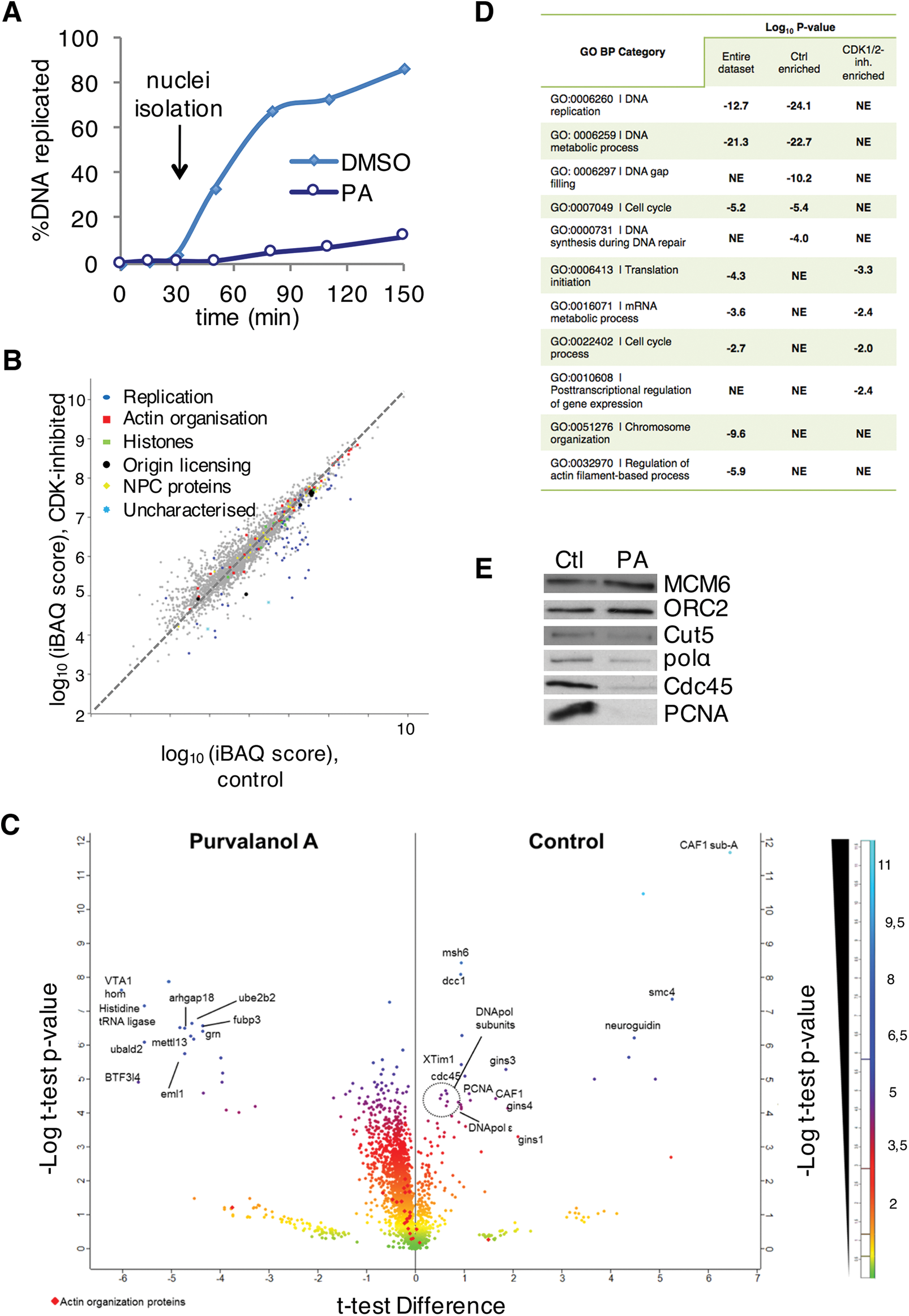
The proteome of replicating nuclei. **A** Replication time-course of sperm chromatin in control and Purvalanol A (PA)-treated egg extracts, with nuclei isolated for MS analysis at 50 min. **B** Graphical representation of the identified proteome with relative quantitation data (mean values from 3 replicates). Full dataset, Table S1. **C** Volcano plot combining the fold-change between control and CDK-inhibited conditions with their log10 *P*-values (Student’s t-test). The most significantly differentially abundant proteins are highlighted. **D** GO analysis using DAVID, showing the most highly enriched GO biological processes in each condition (full GO analysis, Table S2). NE: not enriched. **E** Western blots of chromatin fractions from control and PA-treated nuclei used for MS analysis.

**Figure 2.**
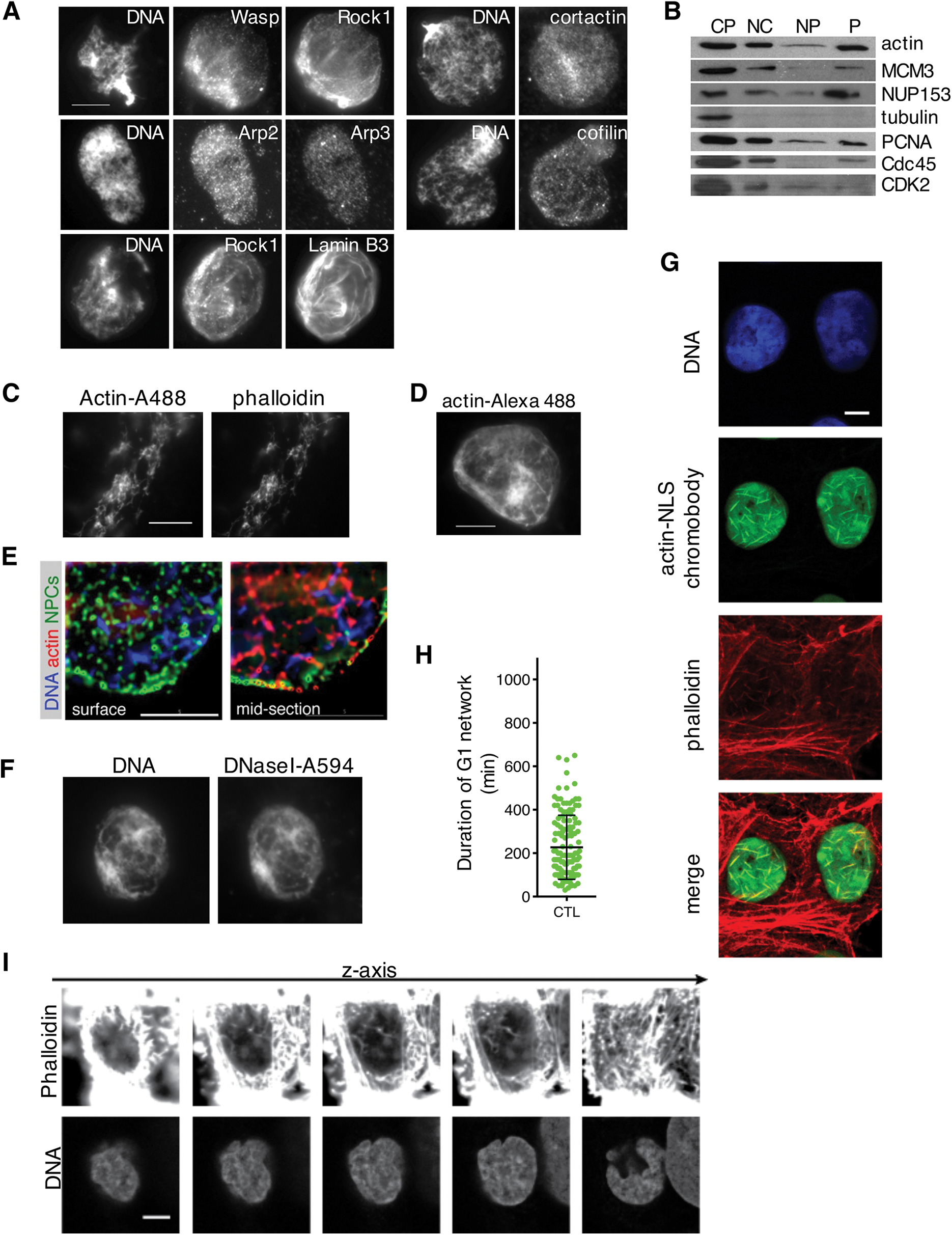
Nuclear actin dynamics during the cell cycle. **A** Immunofluoresence images of the actin regulators indicated, analysed 60 min after sperm head addition. Bar, 10 μm. **B** Western blot analysis of cytoplasm (CP), whole nuclear (NC), nucleoplasmic (NP) and insoluble (P) fraction at 60min-time point during DNA replication, probed with antibodies against proteins indicated. **C** Actin filaments formed by actin-Alexa Fluor 488 in XEE, counterstained with phalloidin. Bar, 10 μm. **D** Immunofluorescence image of nucleus incubated in control extract for 60min in the presence of actin-Alexa Fluor488. Bar, 10 µm. **E** Deconvolved images of 3D optical sections of a nucleus. Left panel, surface; right panel, mid-section. Actin (actin-biotin; red), DNA (blue), NUPs (mAb414, green). Bar, 5 μm. **F** Immunofluorescence images of nucleus formed in control extract, stained at 60 min with DNaseI-Alexa Fluor 594. Bar, 10 μm. **G** Early G1 U2OS cells expressing actin-NLS chromobody co-stained with phalloidin and DAPI (DNA). Bar, 5 μm. **H** Duration of early G1 nuclear actin network (mean ± SD, n=135 cells from 3 independent experiments). **I** Serial confocal planes of an early G1 U2OS cell fixed with glutaraldehyde and stained with phalloidin and DAPI. Bar, 5 μm.

Next, we investigated possible cell cycle regulation of endogenous nuclear actin dynamics in living cells with an actin chromobody (Chromotek®) modified by the addition of a nuclear localisation signal (NLS – see materials and methods). An identical tool was independently developed recently (Plessner *et al*, 2015). We concurrently followed the DNA replication programme using a second chromobody to visualise endogenous PCNA (Burgess *et al*, 2012). Interestingly, we found that a dynamic network of actin filaments formed in most early G1-nuclei (Fig 2G). Filaments disassembled after an average of 200 minutes, in mid-late G1 (Fig 2H; Movie 1). These G1 actin filaments could be stained with phalloidin (Fig 2G), which, importantly, also labelled a G1 nuclear actin network in cells not expressing the chromobody (Fig 2I). Expressing Lifeact-GFP-NLS also revealed nuclear actin filaments (Supplementary Figure 2A). However, this probe disrupted nuclear actin dynamics as although the filaments appeared in G1, they became longer and stable and cells did not divide (Movie 2).

In mouse fibroblasts, formins promote formation of nuclear actin filaments in the serum response (Baarlink *et al*, 2013). Specific formin inhibition with SMIFH2 (Rizvi *et al*, 2009) induced stabilisation of long nuclear actin filaments or patches in the majority of cells (Supplementary Figure 2B, C; Movie 3,4). The former is similar to the effect of formin inhibition on actin dynamics in a reconstituted *in vitro* system (Rizvi *et al*, 2009), and suggests that SMIFH2 stabilised long nuclear actin filaments by preventing formin-mediated nucleation of new filaments, coupled with formin-independent elongation of existing ones.

We subsequently employed XEE to investigate effects of modifying actin dynamics on nuclear actin independently of cytoskeleton-environment interactions and of transcription. First, we used recombinant actin regulatory proteins, as well as different drugs that modify actin dynamics, in preassembled nuclei in XEE (Fig 3A, B). Cytochalasin D (CytD), jasplakinolide, or purified Arp2/3 and GST-WASP-VCA proteins all strongly increased total nuclear actin, which was mostly insoluble. The effects of CytD, which binds the barbed (plus)-end of F-actin, arresting both polymerisation and depolymerisation at the plus end (Schliwa, 1982), were reversed by latrunculin A that potently binds actin monomers, impeding filament assembly. Interestingly, Cyt D caused formation of stable nuclear actin filaments that were visualised with phalloidin (Fig 3C). These data suggest that nuclear actin polymerisation and depolymerisation exist in a dynamic equilibrium in XEE.

**Figure 3.**
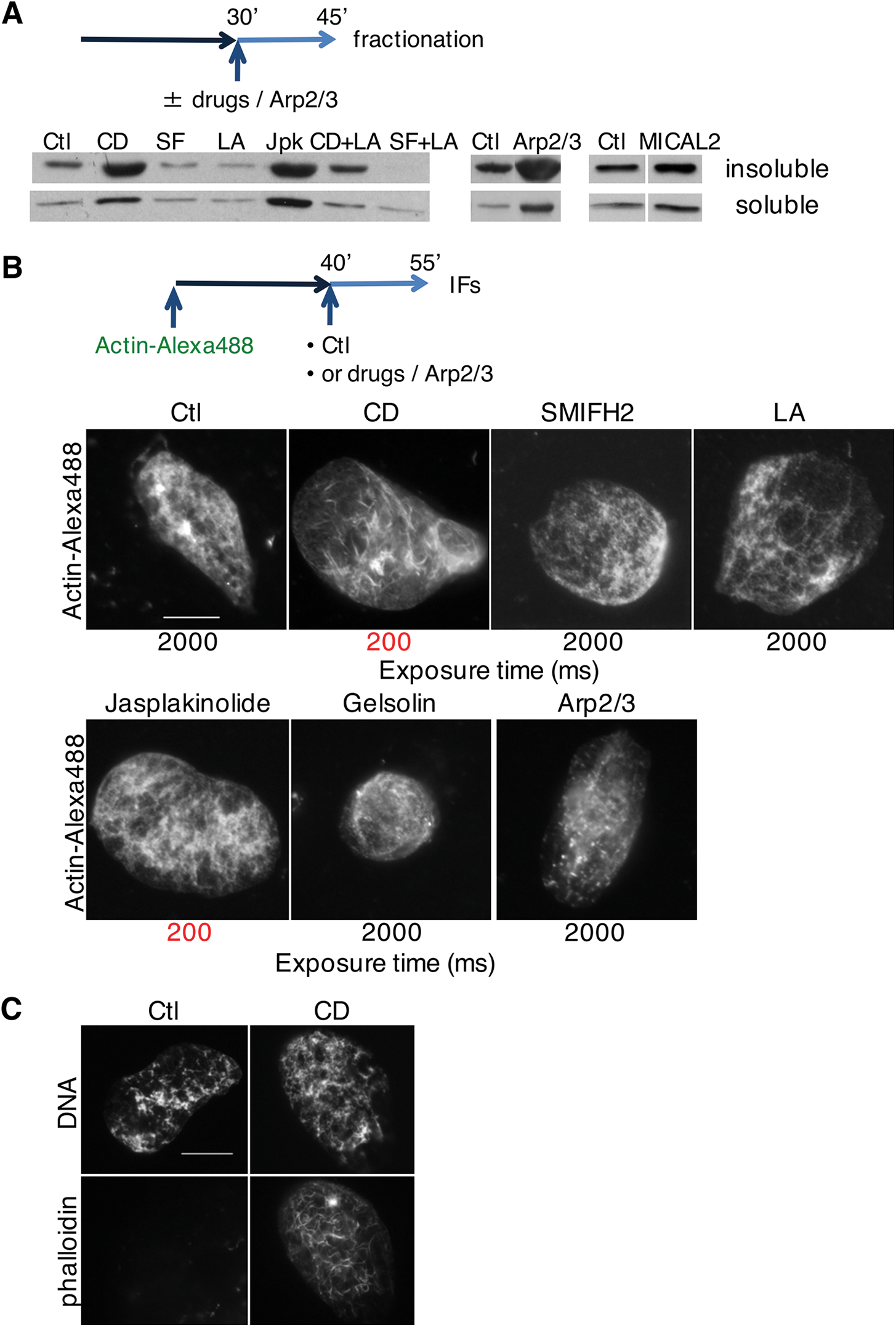
Effect of actin drugs, regulators and probes on nuclear actin dynamics and abundance in *Xenopus* egg extracts. **A** Nuclei were allowed to form for 30 min before drugs (Cyt D, CD; SMIFH2, SF; latrunculin A, LA; jasplakinolide, Jpk; Cyt D and latrunculin A, CD+LA; SMIFH2 and latrunculin A, SF+LA) or Arp2/3 recombinant protein (in combination with VCA domain of WASP) were added, then purified at 45 min. Soluble and insoluble nuclear fractions were blotted for actin. **B** Extract was supplemented with sperm nuclei and actin-Alexa Fluor 488; at 40 min indicated drugs or Arp2/3 and VCA domain of WASP were added, and nuclei were analysed for fluorescent actin at 55 min. Long exposure time (2000ms) was needed to visualise nuclear actin in all conditions with the exception of Cyt D and jasplakinolide (exposure time 200ms, highlighted in red). Bar, 10 μm. **C** Extract was supplemented with sperm nuclei; at 45 min Cyt D (CD) was added and nuclei were analysed at 60 min and stained with phalloidin. Bar, 10 μm.

### Actin dynamics is required for DNA replication

We next assessed the effect of these manipulations of actin dynamics on DNA replication. S-phase entry in G1-synchronised cells was dose-dependently inhibited by SMIFH2 (Fig 4A; Supplementary Figure 3A, B), and PCNA binding to chromatin was reduced (Supplementary Figure 3C). SMIFH2 also abolished general transcription, as determined by 5-ethynyl-uridine (EU) incorporation into newly synthesized RNA (Fig 4B; Supplementary Figure 3D). We then altered endogenous formin activity by expressing GFP-tagged mDia2 diaphanous autoregulatory domain (DAD), either specifically in the nucleus (GFP-DAD.LG.NLS) or cytoplasm (GFP-DAD). Interestingly, neither mDia2-DAD construct interfered with global transcription (Fig 4C). Nuclear, but not cytoplasmic, mDia2-DAD increased the fraction of cells in S-phase (Fig 4D, left), implying that over-activating nuclear formins impedes S-phase progression in a transcription-independent manner. To test whether this might be due to aberrant nuclear actin dynamics, we expressed the nuclear-localised actin mutants S14C and G15S, that favour polymerisation, or the polymerisation-defective R62D) (Supplementary Figure 3E). WT and R62D mutants had no effects on S-phase, but, like formin activation, S14C and G15S mutants increased the fraction of cells in S-phase (Fig 4D, right) and decreased EdU signal intensity (Fig 4E). Thus, promoting nuclear polymeric actin or derepressing formins both impede S-phase progression. Treatment with SMIFH2 of cells expressing PCNA chromobody increased by ten-fold the duration of individual PCNA foci (Fig 4F; Movie 5). Likewise, overexpression of GFP-DAD.LG.NLS immobilised PCNA foci (Movie 6). Impaired PCNA mobility might indicate replication fork stalling, which can generate DNA damage. Indeed, expression of nuclear DAD constructs or SMIFH2 treatment led to formation of DNA double-strand breaks, as shown by the increase in the number of γ-H2AX-positive cells (Supplementary Figure 3F, G). Thus, actin and formin constructs that specifically disrupt nuclear actin dynamics hinder DNA replication.

**Figure 4.**
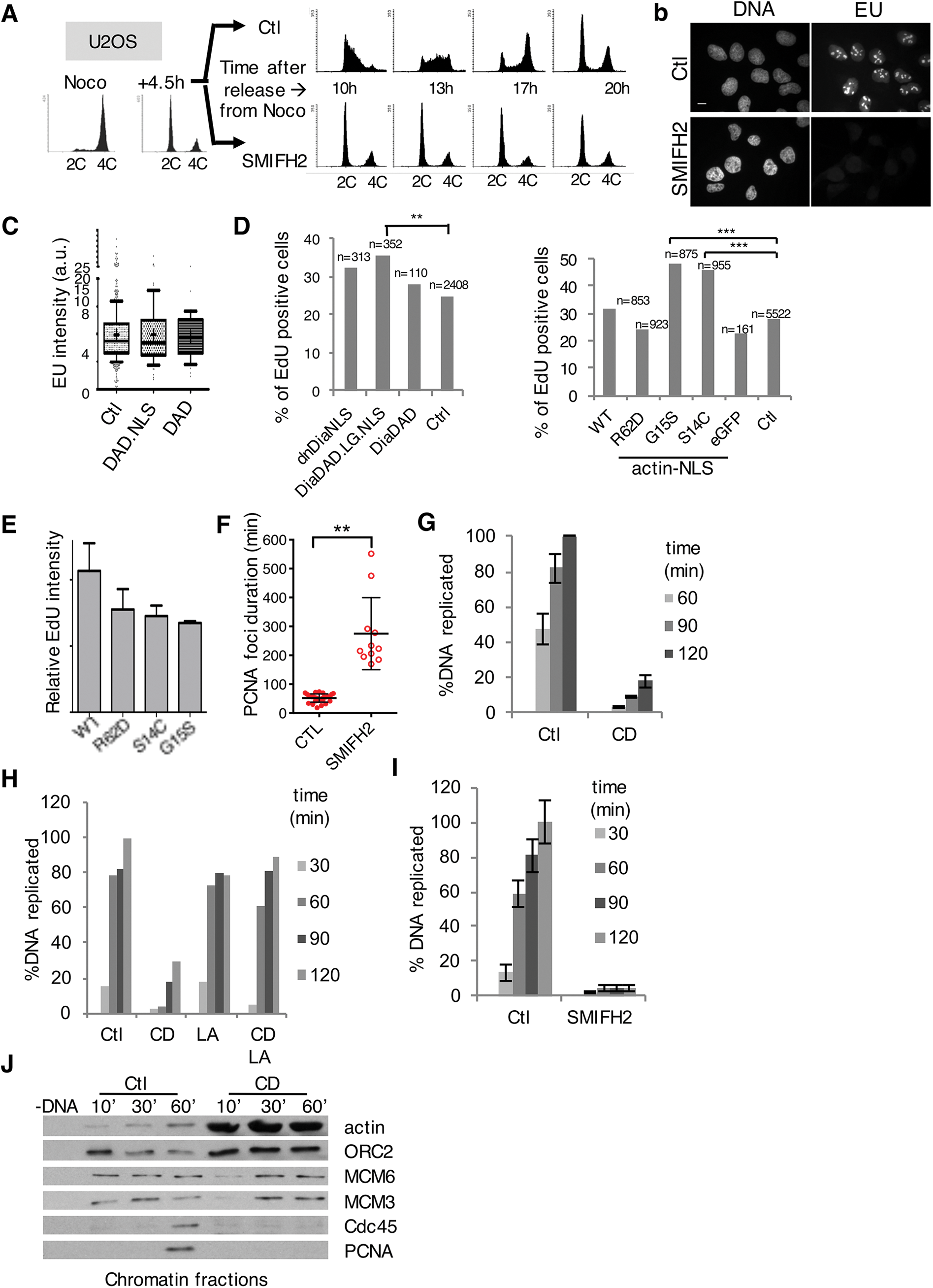
DNA replication requires actin dynamics. **A** FACS analysis of U2OS cells synchronised in G1 and treated with DMSO (Ctl) or SMIFH2 (50μΜ). **B** Immunofluorescent images of control or SMIFH2-treated (1hr pre-treatment, 1hr co-incubation) U2OS cells pulsed for 1hr with EU. Bar, 5 μm. **C** Quantification of EU incorporation (1hr) in U2OS cells, control or transfected with mDia2 DAD.NLS or DAD constructs (n>100, 61 and 47, respectively). **D** Quantification of EdU incorporation from 2 independent experiments after a 60 min-pulse in U2OS cells expressing formin constructs (left) or actin-NLS mutants (right) (**, *p*-value<0.001, ***, *p*-value<0.0001; Student’s t-test). **E** Quantification (mean ± SEM) of EdU intensity from **D** normalised to the non-transfected cells. **F** Duration of PCNA foci in U2OS cells expressing PCNA chromobody (each dot represents the mean of 5-7 foci per cell, mean ± SD of 29 and 11 cells, respectively, per condition from 2 independent experiments). **G** Chromosomal DNA replication determined by ^33^P-dCTP incorporation assay in control conditions or with Cyt D (CD); mean ± SEM of 8 independent experiments. **H** DNA replication assessed in control extract, or extracts supplemented with CytD D (CD), with or without latrunculin A (LA). A representative experiment of 5 independent experiments is shown. **I** DNA replication assays in control (Ctl) or formin-inhibited (SMIFH2; 500M) extract;μmean ± SEM of 8 independent experiments. **J** Chromatin loading of pre-RC and pre-IC factors in control conditions (Ctl) or with Cyt D (CD).

To assess possible effects of altered nuclear actin dynamics on DNA replication independently of transcription, we used XEE. Arresting actin dynamics with CytD inhibited DNA replication (Fig 4G). Combining CytD with jasplakinolide or gelsolin protein was synergistic (Supplementary Figure 4A), whereas latrunculin A, or recombinant cofilin, which severs actin filaments and dissociates monomers, rescued replication (Fig 4H; Supplementary Figure 4B). Inhibiting formins using SMIFH2 or compound 2.4 (Gauvin *et al*, 2009), or inhibiting Arp2/3 with CK-666 (Nolen *et al*, 2009) but not its inactive analogue, CK-689, also blocked DNA replication, as did recombinant MICAL2 protein (Fig 4I; Supplementary Figure 4C–E). Taken together, these results indicate that actin dynamics is required for DNA replication in XEE, independently of transcription. Disrupting actin dynamics inhibited conversion of pre-replication complexes containing ORC and MCMs to pre-initiation complexes (pre-IC) containing Cdc45 and PCNA (Fig 4J; Supplementary Figure 4F). We thus tested possible interactions of endogenous actin with replication factors PCNA, MCMs and RPA by Proximity Ligation Assay (PLA). We could readily detect sites of DNA replication by PLA. Actin interacted with PCNA, but not with RPA or MCMs, indicating that it is not present at replication sites (Supplementary Figure 4G, H). Actin physically associated with PCNA, as confirmed by pulldowns from nuclei using immobilised actin-binding peptide Lifeact (Supplementary Figure 4I).

### Actin dynamics is required for NPC formation and nuclear transport

In XEE, inhibiting actin dynamics prevented DNA decondensation and growth of nuclei (Fig 5A), while nuclei in SMIFH2-treated U2OS cells were smaller and misshapen (Fig 5B). These observations suggested that actin dynamics might be required for nuclear assembly or transport, both of which are essential for DNA replication. We therefore examined nuclear pores in XEE by 3D structured illumination microscopy and whole-mount field-emission scanning electron microscopy (FEISEM). Upon SMIFH2 treatment, nucleoporin (NUP) staining was disorganised, with dense NUP clusters between NUP-free regions (Fig 5C). While most NUPs were present, NUP160 was undetectable, and the levels of NUP107, Gp210, NUP358, NUP214 and NUP183 were decreased (Supplementary Figure 5A). Furthermore, in nuclei formed in the presence of SMIFH2 or CytD, we could not detect nuclear pores with electron microscopy (Fig 5D).

**Figure 5.**
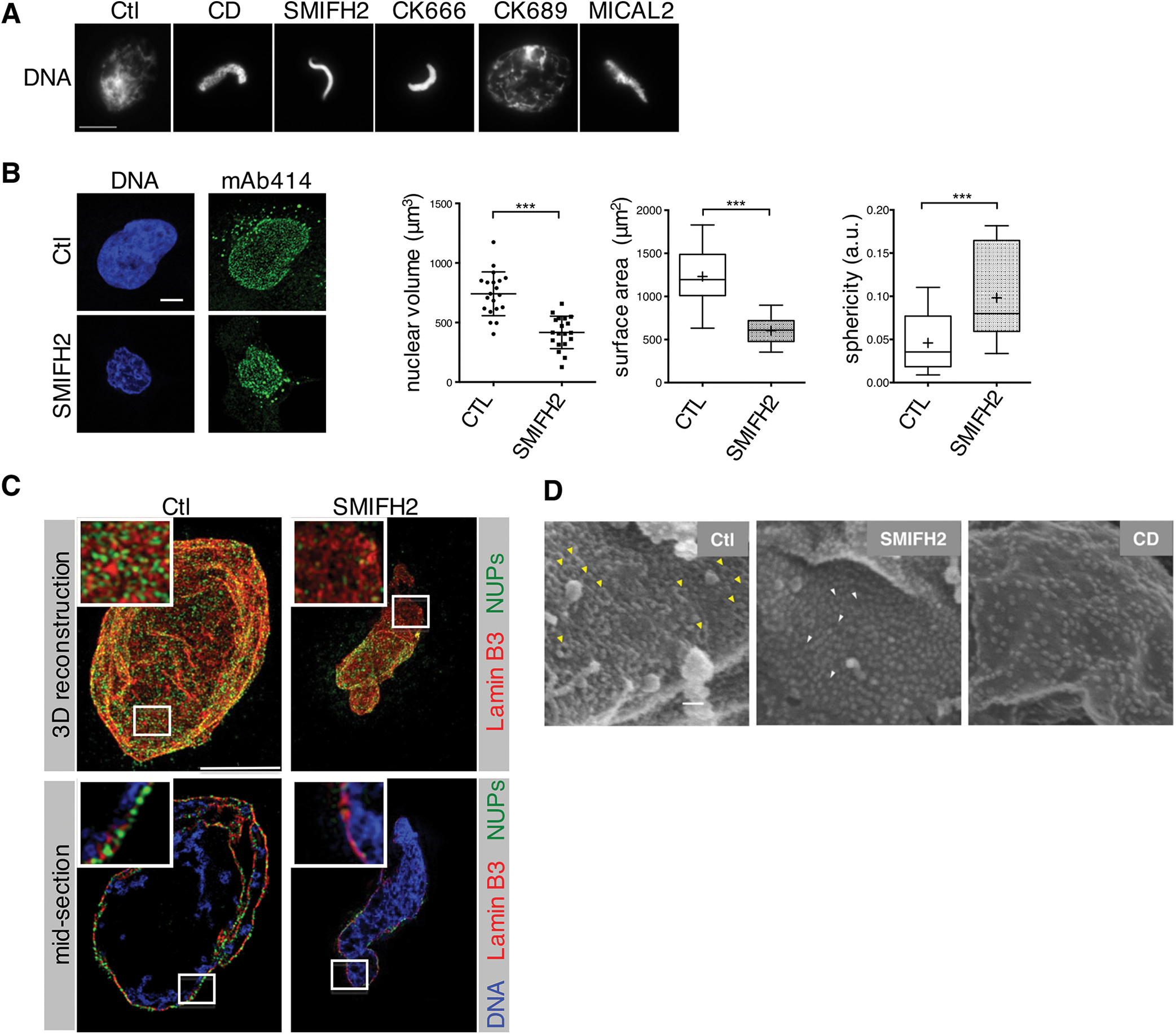
NPC formation requires actin dynamics. **A** Immunofluorescence images of nuclei formed either in control extracts or in the presence of indicated drugs or MICAL2, analysed at 60 min. Bar, 10 μm. **B** Left, confocal planes of nuclei of cells treated with DMSO (Ctl) or SMIFH2 (50)μΜ for 4 h, stained with mAb414 and DAPI (DNA). Bar, 5μm. Right, characterisation of nuclear morphology (mean ± SD of 20 cells from 2 independent experiments; ***, p-value <0.0001, Student’s *t*-test). **C** 3D-SIM images of the nuclear lamina (red), NUPs (mAb414, green), in control or formin-inhibited (SMIFH2) conditions. A reconstructed 3D image (top) and a section (bottom) of the same nucleus are shown. In sections, DNA is shown (blue). Bar, 5 μm. **D** Scanning electron microscopy (FEISEM) images of nuclei formed in the presence of DMSO (Ctl), Cyt D or SMIFH2 at 50min. Representative NPCs (yellow arrowheads) or incompletely formed NPCs (white arrowheads). Magnification x40,000. Bar, 100nm.

We next tested nuclear transport using NLS-tagged GST-GFP. In nuclei formed in CytD- or SMIFH2-treated extracts, the nuclear membrane was intact, but the probe did not accumulate inside nuclei (Fig 6A; Supplementary Figure 5B). Early steps in NPC assembly were unaffected as Elys, importin β, FG-NUPs and RCC1 all bound DNA with similar kinetics to controls (Supplementary Figure 5C). The subsequent step involves RanGTP-mediated release of nucleoporins from importin-*β* (Bai *et al*, 2014). Since the same mechanism also governs active nuclear transport through mature NPCs, we assessed effects of modifying actin dynamics on nuclear transport in preassembled nuclei with NLS-tagged GST-GFP (Fig 6B, scheme). We used wheat germ agglutinin (WGA) that blocks NPC function and importazole that disrupts Ran-importin-*β* interaction (Soderholm *et al*, 2011), and inhibited CDK with PA, which blocks DNA replication without affecting NPC function. As expected, importazole blocked DNA replication (Supplementary Figure 5D). Neither CytD nor SMIFH2 affected integrity of pre-formed NPCs, as assessed probing passive transport with fluorescently labeled dextrans (Mohr *et al*, 2009) (Supplementary Figure 5E). Nevertheless, disrupting actin dynamics abolished nuclear transport, as did WGA and importazole, whereas PA had no effect (Fig 6B).

**Figure 6.**
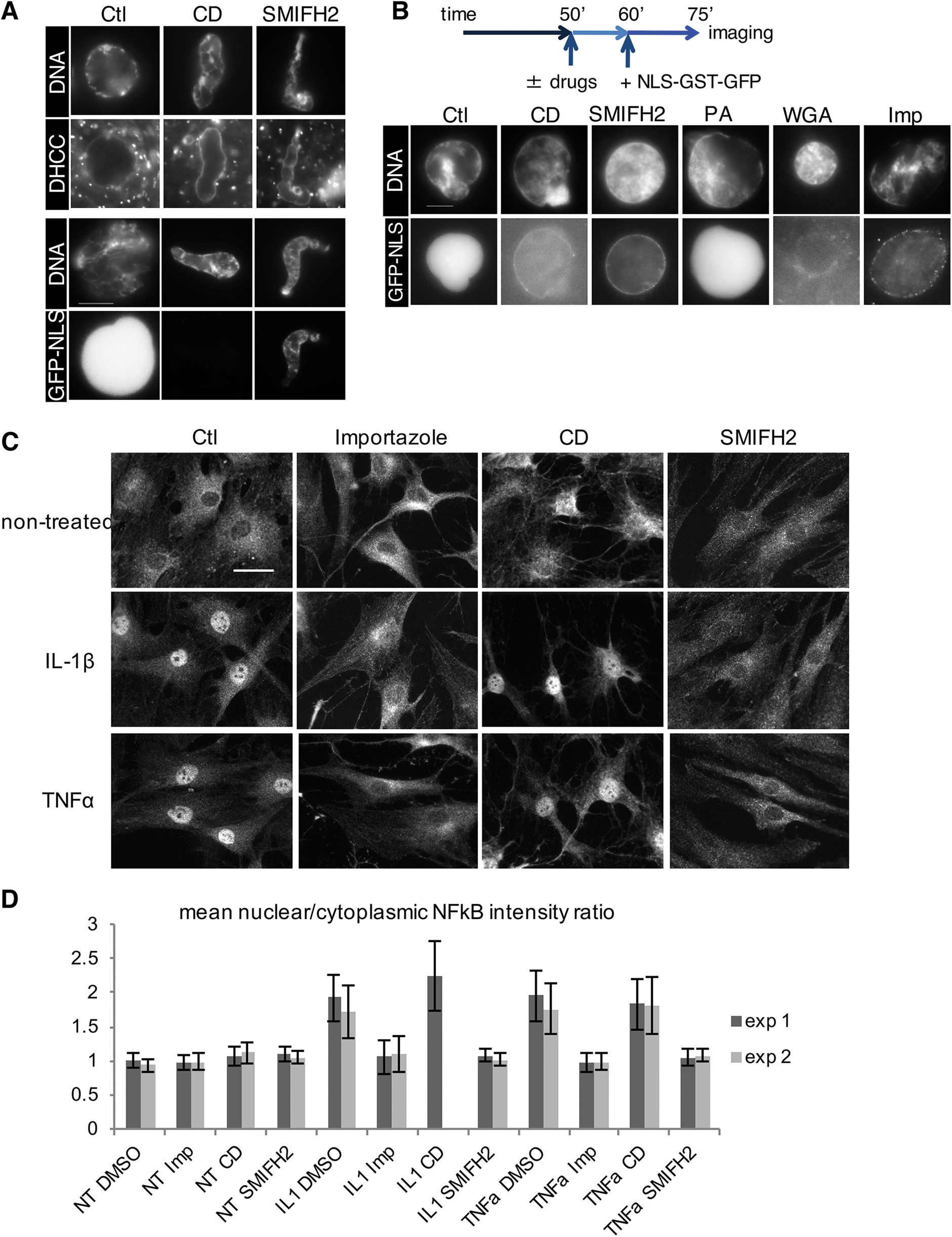
Nuclear transport requires actin dynamics. **A** Immunofluorescence images of nuclei formed in control or Cyt D (CD)- or SMIFH2-treated extracts. Nuclear membranes were visualised with the lipid dye DHCC; nuclear transport was assayed with NLS-tagged GST-GFP protein added at the onset of experiment. Nuclei were analysed at 60 min. Bar, 10 μm. **B** Top, scheme: nuclei were supplemented which Cyt D (CD), SMIFH2, PA, WGA or importazole (Imp) at 50 min; NLS-GST-GFP was added at 60 min and nuclei were imaged at 75 min. Bar, 10 μm. **C** Immunofluorescence images of RA-FLS fibroblasts, treated for one hour with DMSO (Ctl), Importazole (50μM), Cyt D (40μM), or SMIFH2 (50μM), subsequently stimulated or not with IL-1β or TNF-α, stained for NFκB. Bar, 20μm. **D** Quantification of the data presented in **C**. Mean nuclear/cytoplasmic NFκB intensity ratio (±SD) of two independent experiments using two different fibroblast sources; n≥400 for each condition; CytD sample was lost in exp 2.

We next investigated whether actin dynamics is required for importin-dependent nuclear transport in human cells. We analysed nuclear translocation of endogenous NF-κB (Transcription factor p65) in primary human fibroblasts upon stimulation with IL-1β (Interleukin-1 beta) or TNFα (Tumour necrosis factor alpha). Treatment with the cytokines releases NF-κB from its inhibitor IκB (I-kappa-B) and triggers its nuclear translocation, thus bypassing possible effects of cytoplasmic actin disruption on cell shape and NF-κB regulation (Németh *et al*, 2004; Sero *et al*, 2015). This allowed us to study effects of CytD or formin inhibition on NF-κB nuclear translocation itself. Importazole, as expected, strongly reduced NF-κB nuclear translocation. CytD had only a moderate effect, probably because it strongly affected cell shape, which has been reported to alter cytoplasmic NF-κB regulation (Németh *et al*, 2004; Sero *et al*, 2015). In contrast, SMIFH2 did not significantly change cell shape, but almost completely abolished NF-κB nuclear translocation (Fig 6C, D).

### Arresting actin dynamics hinders cargo release from importin

Next, using XEE, we investigated the mechanism whereby inhibiting actin dynamics disrupts nuclear transport. Since FG-NUPs are cargo of importin-β during NPC formation in extracts, we compared FG-NUP-importin interactions in control and CytD- or importazole-treated extracts. Both treatments increased binding of importins to FG-NUPs (Fig 7A). We then analysed effects of CytD and SMIFH2 on binding of importin-*β* to cargo in nuclei. We immunoprecipitated FG-NUPs from nuclear extracts and immunoblotted for PCNA, as well as an unrelated cargo, TPX2 (Targeting protein for Xklp2-A). Importantly, SMIFH2 treatment severely decreased the abundance of all tested proteins in the nuclei. Both treatments resulted in loss of PCNA from nuclei, suggesting that nuclear import of PCNA no longer counterbalances its export. CytD but not SMIFH2 strongly increased the NUP-actin interaction (Fig 7B). PCNA and TPX2 bound similarly to NUPs, indicating that cargo binding is not altered by actin. We then tested whether cargo release in the nucleoplasm was affected. This depends on productive Ran-importin interactions (Lowe *et al*, 2010). Ran binding to importin-β and RCC1, its GTP exchange factor, was not altered by CytD or SMIFH2. However, CytD strongly promoted actin-Ran interaction and TPX2 remained bound to importin-β, suggesting that increased actin binding might hinder cargo release (Fig 7C). TPX2 could not be detected in pull downs from nuclei with SMIFH2, probably as a result of its elimination from the nucleus. Actin binding to Ran could be reconstituted with purified proteins (Fig 7D), and it was independent of whether Ran was in its GDP- or GTP-loaded form (Fig 7E). CytD had no effect on RanGTP levels but promoted nuclear RanGTP-actin binding, and latrunculin A reversed this phenotype (Fig 7F). SMIFH2 marginally increased actin binding to RanGTP, an effect similarly cancelled by latrunculin A. These results suggest that treatments that increase actin binding to RanGTP prevent cargo release from importins, but that this is independent of alterations of nuclear actin levels.

**Figure 7.**
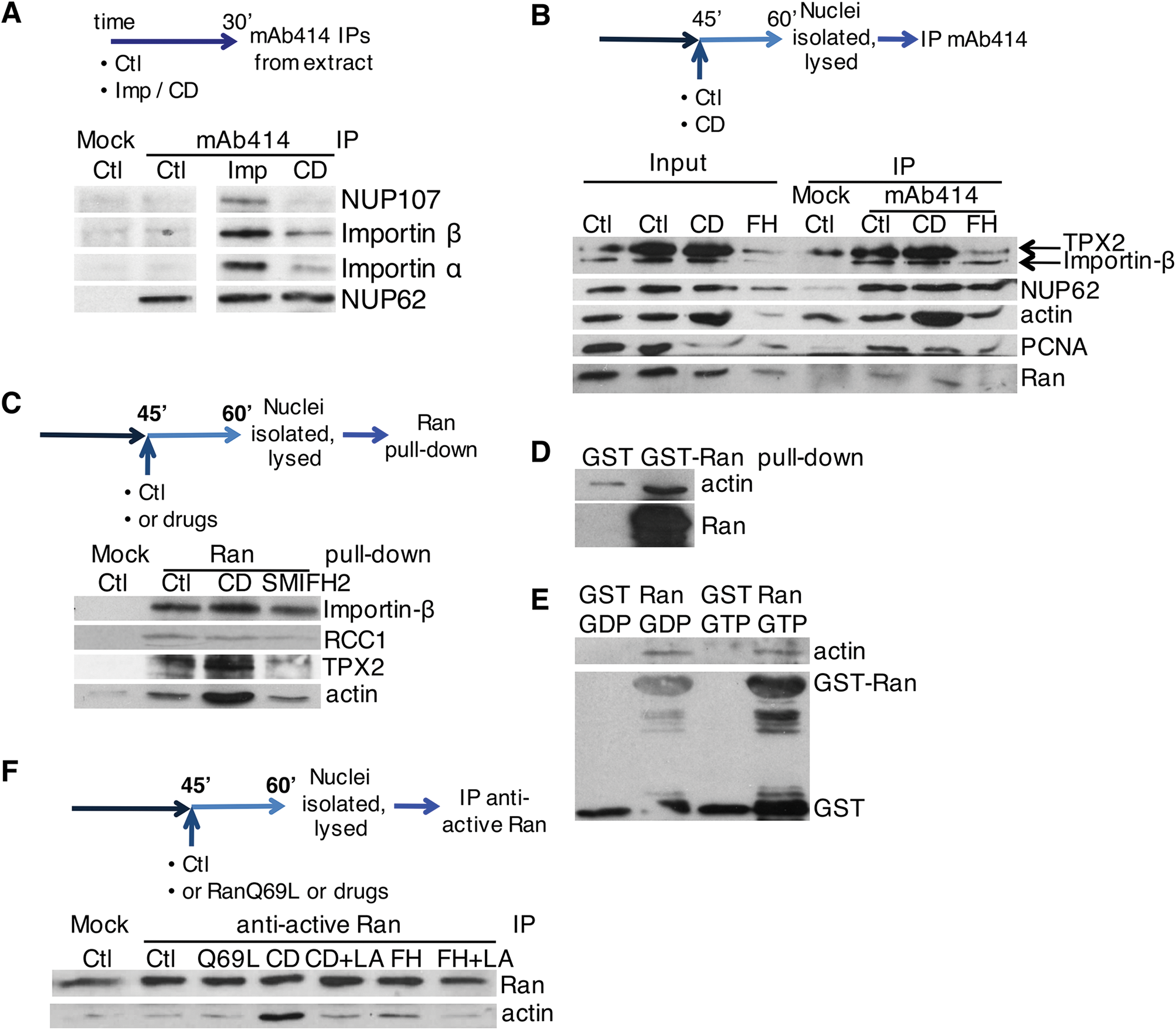
Disruption of actin dynamics hinders cargo release from importin. **A** Scheme: control extract (Ctl) or extract treated with Cyt D (CD) or importazole (Imp) was incubated for 30 min and immunoprecipitated with mAb414 (Mock IP, no antibody added). Beads were blotted for the proteins indicated. **B** Scheme: nuclei, formed in control extract, or with Cyt D (CD) added at 45 min, were purified at 60 min and mAb414 used for immunoprecipitation; 10% of lysed nuclei was used as input. Beads were blotted with the proteins indicated. **C** Scheme: nuclei were incubated in extract supplemented at 45 min with Cyt D (CD) or SMIFH2, lysed and pulled-down with GST-Ran WT immobilised on glutathione beads, or control beads (Mock). Beads were blotted with the proteins indicated. **D** *In vitro* pull-down using GST or GST-RanWT immobilised on glutathione beads and actin-biotin; beads were blotted for actin and Ran. **E** *In vitro* pull-down between actin-biotin and glutathione-immobilized GST or GST-Ran WT, pre-loaded with either GTP or GDP; beads were blotted for actin or GST. **F** Scheme: nuclei, formed in control extract, supplemented at 45 min with RanQ69L, Cyt D (CD) or SMIFH2 (FH), without or with latrunculin A (LA), were lysed and immunoprecipitated with anti-active Ran antibodies (Mock IP, no antibody added). Beads were blotted for actin and Ran.

### Formins act in parallel with CDK to promote pre-IC formation

While nuclear transport is required for nuclear assembly and S-phase onset, our results in human cells indicated that inhibiting formins impairs DNA replication even after nuclear assembly. It was thus important to discriminate whether the roles of formins in DNA replication were exclusively due to their requirement for NPC formation and function. If so, it would suggest that there is a continued requirement for nuclear transport throughout DNA replication. To address this question, we determined execution points for biochemical activities at successive phases of DNA synthesis. We thus used XEE and performed nuclear transfer experiments in which nuclei were isolated from one extract and transferred to another with different conditions. We first used WGA to block existing NPC function, or prevented new NPC formation by depleting nucleoporins with WGA (WGA-bpΔ). As expected, both treatments blocked DNA replication (Supplementary Figure 6A, B). We then combined these treatments with nuclear transfers. Nuclei were formed in an extract where replication licensing was prevented by adding recombinant geminin (Fig 8A, scheme; Supplementary Figure 6C). These nuclei were then transferred into a second extract with added recombinant Cdt1, to release the licensing block, and containing SMIFH2, WGA or vehicle, or depleted of NUPs. In nuclei transferred into NUP-depleted extract, DNA replication occurred, albeit less efficiently (Fig 8A, WGA-bp∆). However, replication was totally blocked by the further addition of SMIFH2 (Fig8A, WGA-bp∆ +SMIFH2). Blocking existing NPCs with WGA also prevented replication (Fig 8A, +WGA). Therefore, once nuclei have been correctly formed, DNA replication can initiate in the absence of further NPC formation, but not if formins or existing NPCs are inhibited.

**Figure 8.**
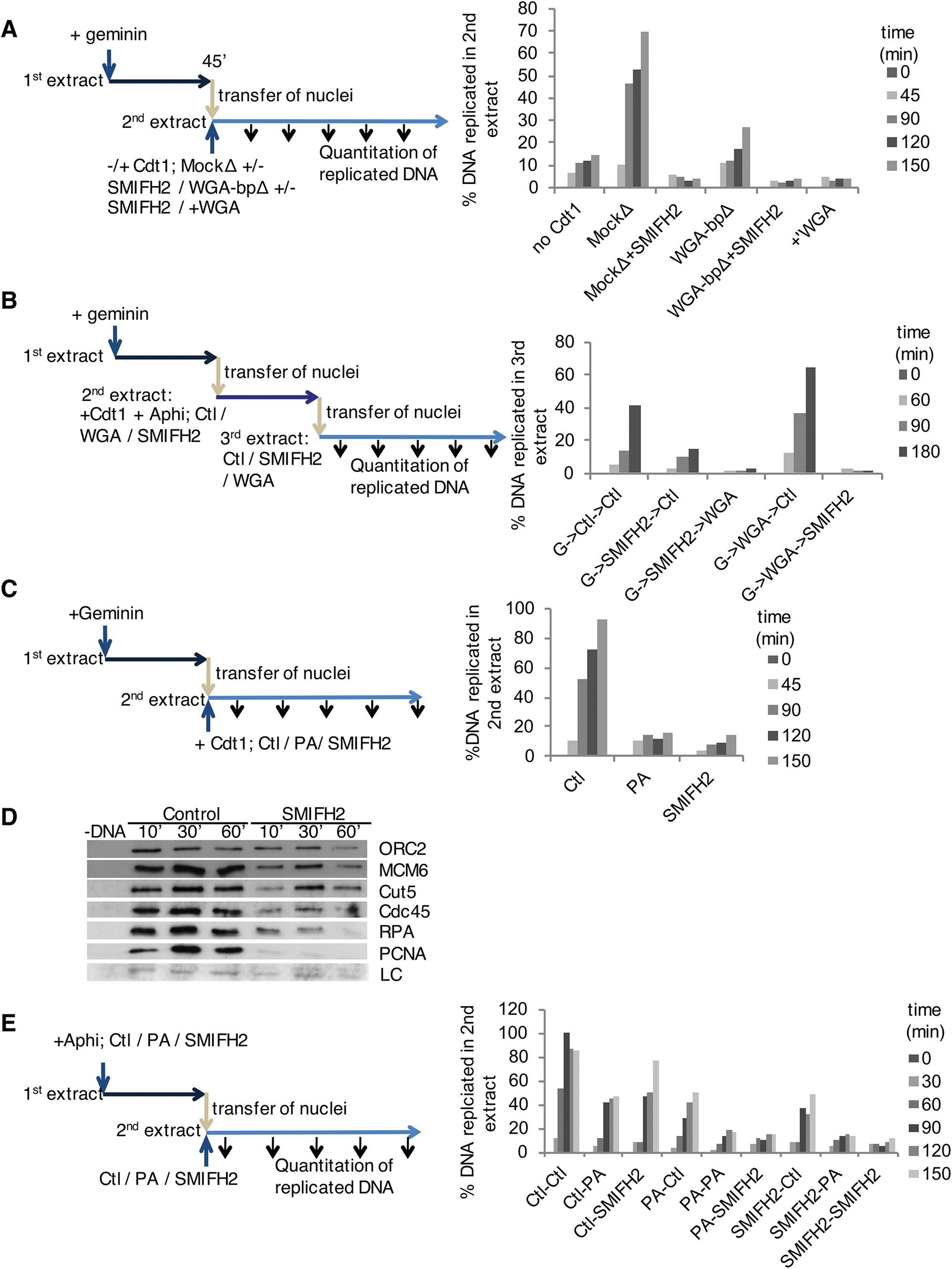
Formins promote pre-IC formation in parallel with CDK. **A** Scheme: nuclei were formed in the first extract containing geminin, then transferred to a second extract, with Cdt1, which was either Mock- (MockΔ) or WGA-binding protein- (WGA-bpΔ) depleted, with or without addition of SMIFH2; or where WGA was added (+WGA). Replication efficiency was measured in the second extract. **B** Scheme: double reciprocal nuclear transfer, from geminin-treated extract into either SMIFH2- or WGA-treated Cdt1-containing extract, with aphidicolin; and then into a third extract with the alternative condition. DNA replication was assessed in the third extract. **C** Scheme: nuclear transfer experiment, in which first extracts contained geminin; second extracts contained Cdt1 and were controls (Ctl), or CDKs (PA) or formins (SMIFH2) were inhibited. DNA replication was assessed in the second extract. **D** Chromatin was purified from second extracts of experiment in **c** and blotted for the proteins indicated. LC, loading control. **E** Scheme: reciprocal nuclear transfer experiment, in which first extracts contained aphidicolin (Aphi) and either PA or SMIFH2 or were controls; nuclei were isolated from each extract after 45 min and transferred to the same combination of conditions. DNA replication was assessed in the second extract.

This suggests that formins might have a role in the continued function of NPCs in DNA replication. To further investigate such a possibility, we used a double reciprocal nuclear transfer. First extracts contained geminin and second extracts contained either SMIFH2 or WGA. Nuclei were further transferred into a third extract with the alternative condition or to a control extract (scheme, Fig 8B). Neither transfer from WGA-into-SMIFH2, nor from SMIFH2-into-WGA allowed DNA replication in the third extract (Fig 8B). Therefore, ongoing nuclear transport and formin activity are required to promote DNA replication in fully formed nuclei.

Given that CDK activity is essential for pre-IC formation, but not for nuclear assembly or transport, we next performed reciprocal nuclear transfer between CDK-inhibited (PA) and WGA-treated extracts. Replication was abolished when nuclei were transferred from PA to WGA (Supplementary Figure 6D), confirming that active nuclear transport is required in parallel with CDK to promote pre-IC formation. Assuming formin function is to allow nuclear transport, formin activity should therefore be essential for pre-IC formation. We tested this by transferring nuclei from geminin-containing extract to a second extract, treated with either PA, SMIFH2 or vehicle, and quantifying DNA replication. As expected, both PA and SMIFH2 prevented replication in preassembled nuclei (Fig 8C). Chromatin-bound PCNA was essentially undetectable in SMIFH2-treated nuclei (Fig 8D), showing that formin activity is required after nuclear assembly in XEE to allow pre-IC formation and DNA replication, as in somatic cells. We therefore next determined whether formins and CDK act sequentially or in parallel. We performed reciprocal nuclear transfer experiments between a formin-inhibited and a CDK-inhibited (PA) extracts (Fig 8E). First extracts additionally contained aphidicolin to prevent replication fork progression, so that replication in the second extract reflected pre-IC assembly. DNA replicated when transferred from a control first extract to a second containing SMIFH2 or PA, but not when transferred from PA to SMIFH2, nor, as expected, when transferred from SMIFH2 to PA (Fig 8E). Therefore, formins are required in parallel with CDK to promote pre-IC formation.

### Nuclear formin activity controls chromatin loading of PCNA and CDKs

Finally, we tested whether nuclear formins might have roles in DNA replication that are independent of nuclear transport. To do this, we performed a nuclear transfer experiment where first extracts contained leptomycin B to inhibit the exportin Crm1, allowing nuclear accumulation of replication factors, and PA to inhibit initiation of replication. Second extracts contained SMIFH2 or vehicle, with or without leptomycin B (Fig 9A). DNA replicated efficiently in control second extracts. Without leptomycin B in the second extract, SMIFH2 treatment decreased nuclear levels of both CDKs and PCNA, explaining why continuous nuclear transport is required for efficient DNA replication. Leptomycin rescued CDK and PCNA levels, but DNA still could not replicate (Fig 9B), and neither CDK nor PCNA were present on the chromatin (Fig 9C). Therefore, nuclear formin activity is further required for loading of pre-IC components onto chromatin. Decreased loading of PCNA would lead to reduced origin firing and fork stalling, which would explain impaired S-phase progression observed in somatic cells and induction of DSBs (Fig 4D–F; Supplementary Figure 2A–C, F, G).

**Figure 9.**
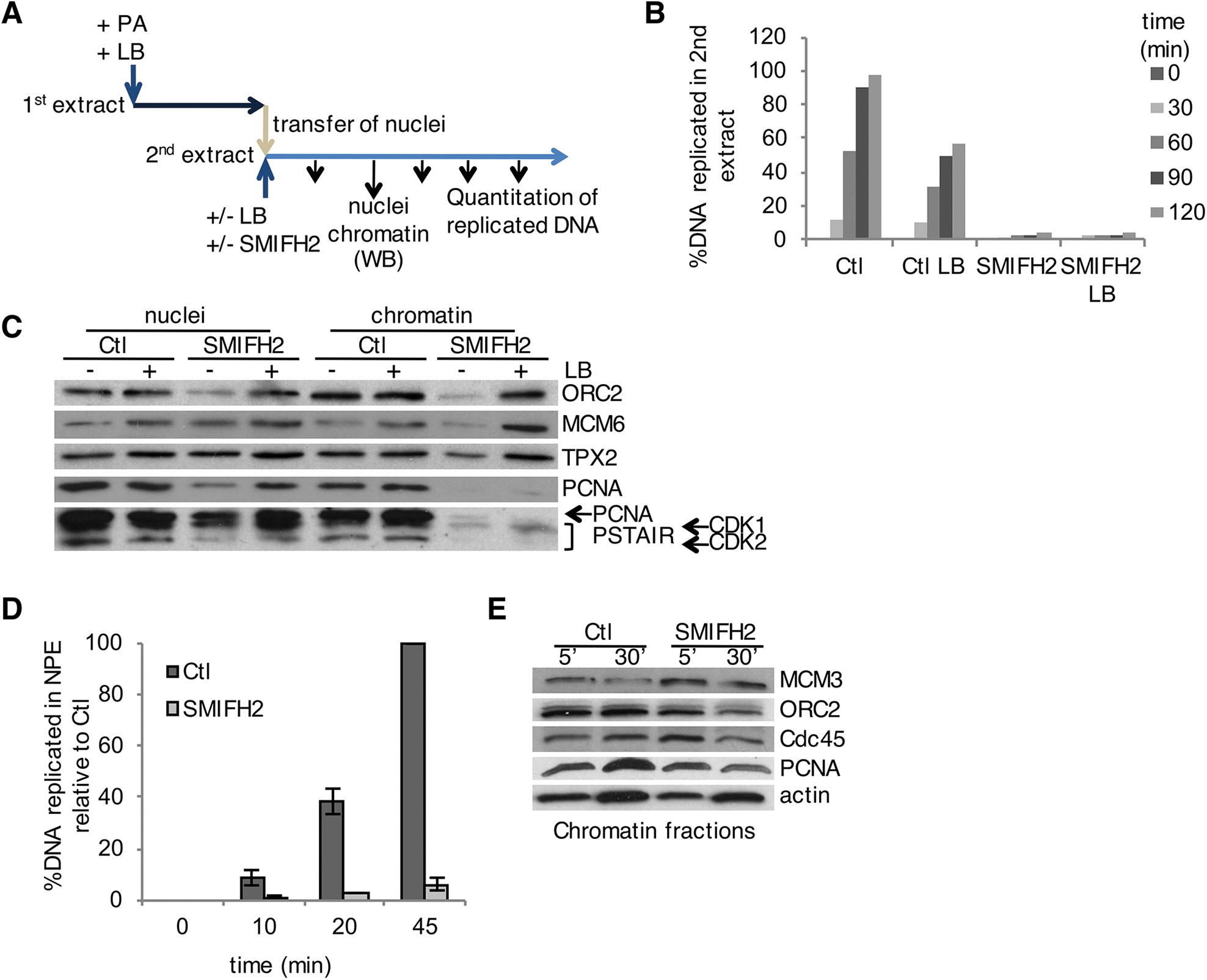
Nuclear formin activity controls chromatin loading of PCNA and CDKs. **A-C** Scheme: nuclear transfer from the first extract, where active nuclear export (Leptomycin B, LB) and CDK (PA) were inhibited, into the second extract, either control or treated with SMIFH2, containing leptomycin B or not. **B** DNA replication was assessed in the second extract. **C** Total nuclear and chromatin-associated replication factors in nuclei isolated at 60 min were analysed by Western blotting. **D** Chromosomal DNA replication in NPE determined by ^33^P-dCTP incorporation assay in control conditions (Ctl) or in the presence of SMIFH2; mean ± SEM of 4 independent experiments. **E** Chromatin loading at 30 min of indicated replication factors in NPE, in control (Ctl) or SMIFH2-treated extracts.

We surmised that chromatin loading of PCNA and CDK, as well as DNA replication, might not require formins in nucleoplasmic extracts (NPE). They are highly concentrated and DNA can replicate in the absence of a nuclear envelope (Walter *et al*, 1998). DNA in control NPE replicated efficiently, but replication was totally abolished when formin activity was inhibited with SMIFH2 (Fig 9D). A similar effect was observed with the 2.4 formin inhibitor (Supplementary Figure 6E). Importantly, the initial loading of PCNA and other pre-IC components onto chromatin occurred, but did not increase following initiation of DNA replication as in the control extract (Fig 9E). Thus, formin inhibition specifically prevents chromatin loading of replication components in nuclei, and reveals an additional downstream formin-dependent step in the initiation of DNA replication.

## Discussion

Our study identifies new roles for actin dynamics and formins in controlling cell proliferation. There are several reasons why this might not previously have been observed. First, cell anchorage and the cytoskeleton are involved in growth factor-dependent transcription, *e.g.* of cyclin D1 in mammalian cells (Assoian & Zhu, 1997), as well as degradation of the CDK inhibitor CDKN1A (Densham *et al*, 2009). This is at least partly due to cytoplasmic MST kinase activation and signaling to JNK (Densham *et al*, 2009), obscuring possible effects of altered nuclear actin dynamics. We and others (Serebryannyy *et al*, 2016) find that treatments that induce nuclear actin filaments eliminate global transcription. We show here that manipulating nuclear actin also arrests DNA replication in a transcription-independent manner in somatic mammalian cells. Additionally, we found that specifically interfering with activity of nuclear formins in S-phase and actin dynamics disrupts DNA replication in transcriptionally silent XEE. Second, actin is required in other cell cycle phases, for cortical reorganisation and contractile ring formation (Schroeder, 1973), centrosome separation and mitotic spindle formation (Uzbekov *et al*, 2002; Rosenblatt *et al*, 2004). Thus, genetic mutation or knockdown of actin regulators, which cannot be induced specifically in S-phase, disrupt cell division. Identification of roles for actin dynamics in S-phase can only be achieved using chemical modulation in synchronised cells, or using a system such as XEE where S-phase can be studied independently of other cell cycle phases, transcription and the cytoskeleton, is synchronous and can be broken down into individual steps.

Because XEE are highly concentrated compared to cell culture medium, and there is no active drug transport, far higher concentrations of pharmacological inhibitors are required than in cultured cells. XEE contain around 50 mg/ml protein, of which 5-10% is actin. Thus, there is at least 100 μM actin in extracts. Since effective drug concentrations depend on adsorption, distribution and metabolism, it is expected that several hundred-micromolar concentration of actin drugs is required to elicit phenotypic effects in this system. In contrast, in cells, due to active import, drugs can routinely attain 1000-fold higher concentrations than in the medium (Martinez Molina *et al*, 2013).

Nucleocytoplasmic shuttling of actin (Dopie *et al*, 2012) means that drug effects on cytoplasmic actin have knock-on effects on nuclear actin levels. Indeed, CytD and jasplakinolide both greatly increased nuclear actin levels, and CytD promoted nuclear actin filament stabilisation in XEE. However, adding recombinant GST-WASP-VCA and Arp2/3 also increased nuclear actin but did not promote similar filament formation nor affect nuclear transport and DNA replication. Conversely, formin inhibition did not raise nuclear actin levels nor trigger nuclear actin filament stabilisation, yet inhibited both nuclear transport and DNA replication. Furthermore, addition of purified proteins MICAL2 or gelsolin was inhibitory for replication, while addition of recombinant cofilin could rescue the effects of CytD. Finally, hyperactivation of nuclear formins inhibited S-phase progression. These results imply that deregulated nuclear actin dynamics, rather than an increase in nuclear actin levels or filament formation *per se*, prevents DNA replication.

Our experiments in XEE demonstrate that actin dynamics is essential both for NPC assembly and for nuclear transport, and we reveal one underlying mechanism. Both of these processes involve importin-α/β-mediated cargo binding and subsequent release, which is dependent on the interaction with Ran. Arresting actin dynamics results in increased actin binding to RanGTP, preventing cargo release from importin-β. A similar phenotype has been observed with importazole, which alters Ran-importin interactions without preventing their binding (Soderholm *et al*, 2011). Similarly, the K37D/K152A Ran mutation affects importin-β-Ran interactions, impeding cargo release (Lee *et al*, 2005). Future studies will be required to map the exact interaction sites and determine conformational changes induced by actin binding to Ran, and how this modifies Ran-importin interactions.

Further work will also be required to define whether the observed effects of altering formin activity can be entirely attributed to changes in actin dynamics. We find that activating endogenous nuclear formins in somatic cells, or favouring nuclear actin polymerisation by expressing NLS-tagged actin mutants, arrests ongoing DNA replication. We also find that formin activity is required for DNA replication downstream of nuclear assembly and independently of nuclear transport. In XEE it is required for loading CDKs and PCNA onto chromatin, while in NPE it directly promotes DNA replication. These results suggest that nuclear organisation is not simply required to concentrate replication factors, as assumed from DNA replication in nuclear envelope-free *Xenopus* nucleoplasmic extracts (Walter *et al*, 1998). It will be important to define the precise mechanism of these formin-dependent steps in DNA replication.

In conclusion, together with accumulating evidence for important roles in chromatin regulation and transcription, our study strongly reinforces the notion that actin dynamics and formins have critical effects on essential nuclear processes.

## Materials and Methods

### Antibodies

Antibodies used are as follows: XCdc45, XCdc6, XRPA, XMCM3, XCut5 (gifts from M. Méchali); XORC2, XMCM6 (gifts from J. Maller); XCut5 (gift from D. Maiorano); PCNA (Abcam; ab18197, or Oncogene Science NA03); PSTAIR (Sigma-Aldrich; P7962); human Cdc6 (H-304, Santa Cruz Biotechnology; SC-8341); actin (Sigma-Aldrich, clones A2066 or AC-15; Hypermol, clone 2G2); Ran (Santa Cruz Biotechnology, C29; SC-1156); active Ran (NewEast Bioscences; 26915); cofilin (Abcam; ab42824); Arp2 (Abcam; ab47654); mDia2 (One World Lab; 11016); NFκB p65 (A) (Santa Cruz Biotechnology; SC-109); XNUP107, XNUP62, XNUP153 (gifts from B. Heulsmann); Elys (gift from J. Blow); γH2A.X pSer129 (Millipore, clone JBW301); TPX2, XRCC1, HS importin β. In house rabbit polyclonal antibodies against His-tagged *Xenopus* importin α were raised and affinity purified. The original construct for His-tagged human importin-α was a gift of D. Goerlich; XLaminB3 (gift from B. Goldman); WASP (Abcam; ab74904); mAb414 (Abcam; ab50008); ROCK1 (Abcam; ab58305); Arp3 (Abcam; ab49671); Cortactin (Millipore; clone 4F11); tubulin (Santa Cruz Biotechnology; SC-9104); GST (Pierce; MA4-004); biotin (Cell Signaling; D5A7); digoxigenin (Roche, clone 1.71.256).

### Plasmids

The PCNA-TagRFP and actin-TagGFP chromobodies were purchased from Chromotek **®**. To allow endogenous nuclear actin detection, the SV40 nuclear localisation sequence (NLS, ccgcctaagaaaaagcggaaggtg) was added at the C-term of the actin chromobody, or in between the actin Vhh sequence and the TagGFP. The former is essentially identical to the nAC recently published (Plessner *et al*, 2015) but with a different stop codon. Both of our nuclear actin chromobodies gave identical results but only the former was used in this study. The actin-NLS R62D mutant (Baarlink *et al*, 2013) was used as template to generate the actin-NLS WT form and that was subsequently mutated to S14C or G15S. Formin mutants, mDia2-DAD constructs, actin-NLS R62D, and Lifeact-GFP-NLS were gifts from R. Grosse.

### *Xenopus* egg extracts and replication reactions

Interphase egg extracts, chromatin isolation and replication assays were prepared and performed essentially as described (Blow & Laskey, 1986), with minor modifications. In brief, eggs laid overnight in 150mM NaCl were dejellied in degellying buffer (29mM Tris pH 8.5, 110mM NaCl, 5mM DTT); rinsed several times in High Salt Barths solution (15mM Tris pH 7.6, 110mM NaCl, 2mM KCl, 1mM MgSO4, 0.5mM Na2HPO4, 2mM NaHCO3), twice in MMR (5mM HEPES-KOH pH 7.6, 100mM NaCl, 2mM KCl, 0.1mM EDTA, 1mM MgCl2, 2mM CaCl2), before activation with 0.3μg/ml calcimycin ionophore in MMR. Subsequently, two rinses in MMR and two more in SB (50mM HEPES-KOH pH 7.6, 50mM KCl, 2.5mM MgCl2, 5% Sucrose, 0.014% β-mercaptoethanol) followed, while during the last rinse the eggs were transferred on ice and SB was supplemented with protease inhibitors (10μg/ml leupeptin, pepstatin and aprotinin). Eggs were spun down at 200g for 1min and excess of buffer removed before being centrifuged at 16,000g, 4°C for 10 minutes. Protease inhibitors and 10μg/ml cytochalasin B were added to the cytoplasmic fraction. This concentration of cytochalasin B, a much weaker actin drug than cytochalasin D, is required to reduce the viscosity sufficiently that extracts can be obtained by centrifugation but has no effect on DNA replication and does not provoke nuclear actin stabilisation. Extracts were further centrifuged in SW55Ti rotor for 20min at 20k rpm (48,000g) at 4°C. The cytoplasmic layer was extracted with a large-bore needle and syringe, and supplemented with glycerol 3% and ATP regenerating system (10mM creatine phosphate, 10μg/ml creatine kinase, 1mM ATP, 1mM MgCl2) added from a 20x stock. Aliquots were frozen in liquid nitrogen. Where indicated, a 1:100 dilution of cytochalasin D (at final concentration of 400 μM, unless otherwise indicated; Enzo); SMIFH2 (500 μM, unless otherwise stated; Calbiochem); Purvalanol A (200 μM; Sigma-Aldrich); latrunculin A (100 μM, unless otherwise indicated; Enzo); jasplakinolide (100 μM; Enzo); importazole (500 μM, unless otherwise indicated; Sigma-Aldrich); 2.4 formin inhibitor (at indicated concentrations; K216-0385, ChemDiv); CK-666 or CK-689 (Calbiochem), or DMSO solvent only was added to the *Xenopus* egg extracts. Where indicated, extract was supplemented with: recombinant geminin and Cdt1 (40nM; gift from M. Lutzmann); recombinant MICAL2 (48ng/μl of extract; gift from V.N. Gladyshev); WGA (0.2 mg/ml; Calbiochem); aphidicolin (25 μg/ml; Sigma-Aldrich); recombinant cofilin (5 μM; Hypermol; 8419-01); recombinant Arp2/3 complex and GST-VCA (200nM; Hypermol;84101 and 8416-01, respectively); recombinant His-Ran WT and Q96L (used at 5μM; purified as described previously: (Bompard et al., 2005); gelsolin (80ng/μl; Sigma-Aldrich, G8032); dextran-Alexa Fluor 488 10,000MW, and dextran-Rhodamine B 70,000MW (used at 2.5μl/μl; Life Technologies, D-22910 and D-1841). For mass spectrometry analysis, sperm heads were added at concentration of 2800/μl and the insoluble fraction of nucleoskeleton and chromatin was isolated at 50 min from 1ml of extract per condition. Nucleoplasmic extracts (NPE) preparation, analysis of DNA replication efficiency and chromatin loading of replication factors were performed as described. *Chromosomal DNA replication in a soluble cell-free system derived from Xenopus eggs*, AV Tutter and JC Walter, in *Xenopus Protocols. Cell Biology and Signal Transduction*. Humana Press, Totowa, New Jersey 2006). The NPE was supplemented with DMSO, SMIFH2 or 2.4 compound at 1.6% (SMIFH2 final concentration 800μM, unless otherwise stated).

### Cell culture

Cells (U2OS or HeLa) were cultured in DMEM Glutamax (Invitrogen) supplemented with 10% heat inactivated Fetal Bovine Serum (FBS; Invitrogen) and 1x antibiotic mixture (complete medium). Cell lines were tested for mycoplasma contamination regularly. For cell cycle synchronisation, cells were incubated in complete medium containing 2 mM thymidine for 14-16 h. After an 8-10 h release in complete medium, 2 mM thymidine was added again for 20 h. Cells were released and 6-7 h later nocodazole (50-100 ng/mL) was added for additional 5 h. Cells were washed with PBS and complete medium was added for 4.5 h, at which time DMSO or SMIFH2 (50 μM) was added. At indicated time-points, cells were detached by trypsinisation, washed with ice-cold PBS and pellets were collected for FACS and/or immunoblotting analysis. For transient transfections of plasmid DNA, jetPEI or Lipofectamine 2000 or 3000 was used, according to the manufacturer’s instructions (Polyplus Transfection or Invitrogen, respectively).

U2OS cells stabely expressing the nuclear actin or PCNA chromobody were obtained upon Lipofectamine-2000 transfection and selection with 2 μg/ml pyromycin. Clones were obtained by serial dilution.

To analyse NFκB translocation, RA-FLS (rheumatoid arthritis, fibroblast-like synovicites) were prepared as described (J. Morel et al. JBC 2005; 280: 15709-18) (Morel *et al*, 2005). Cells were seeded at 10 000 per well on coverslips in 12-well plates in RPMI medium/5% FBS, allowed to adhere for 24hrs, then starved overnight in RPMI/1% FBS. The following day, fresh medium/1% FBS was supplemented with 0.1% DMSO, importazole (50μM) or SMIFH2 (50μM) for one hour, followed by stimulation with IL-1β (10ng/ml final; Miltenyi Biotec) or TNF-α (10ng/ml final; Miltenyi Biotec) for 30 min. Cells were then washed in PBS, fixed in 3.7% formaldehyde/PBS and proceeded for NFκB immunostaining.

### Immunoprecipitations, pull-downs and nuclear transfers

Glutathione-immoblised GST-Ran wild-type and Q69L mutant were produced as previously described (Bompard *et al*, 2010). For IPs, 10μl of beads (glutathione-Sepharose (GE Healthcare) beads were used as Mock) were washed in PBS and incubated with lysed nuclei (corresponding to 25μl of extract, lysed at 55min; drugs were added at 40min) for 2h at 4°C, washed in 150mM NaCl/PBS, resuspended in Laemmli buffer and analysed by Western-blotting.

For mAb414 and importin-β IPs from egg extract, 10µl of DynaBeads (for mAb414) or 10µl packed protein G-agarose (Roche) beads (for importin-β) were washed with PBS and incubated with antibody for 2h at 4°C, subsequently washed in PBS and incubated with 25µl extract diluted with SB buffer for 2h at 4°C. Beads were then processed as above.

For anti-active Ran IP, 10µl packed protein G-agarose (Roche) beads were incubated with 1µg of antibody for 2h at 4°C, blocked in 10mg/ml BSA/PBS, washed in PBS and incubated with lysed nuclei (corresponding to 25μl of extract) for 2h at 4°C, washed in 0.1% Triton-X 100 / 150mM NaCl / PBS, resuspended in Laemmli buffer and analysed by Western blotting. For actin-Ran *in vitro* pull-down, glutathione-Sepharose (GE Healthcare) beads were pre-incubated with recombinant GST protein (Bompard et al. 2010); 10μl of glutathione-GST and glutathione-GST-Ran beads were washed and blocked in 10mg/ml BSA/PBS, washed in PBS and incubated with 1μg of actin-biotin (Cytoskeleton) for 2h at 4°C, then proceeded as above. For Lifeact-NLS-actin pull-down, 10μl packed Streptavidin-Agarose (Novagen) beads were incubated with 5 nmol Lifeact-NLS-biotin peptide (MG-VADLIKKFESISKEEGDPP-VATPPKKKRK-V-biotin; synthesised by Cambridge Research Biochemicals) for 2h at 4°C, washed in PBS and incubated with lysed sonicated nuclei (corresponding to 40μl of extract) for 2h at 4°C, washed in 150mM NaCl/PBS, resuspended in Laemmli buffer and analysed by Western blotting.

GTP/GDP nucleotide exchange assay with glutathione-immoblised recombinant GST-Ran was performed as previously described (Bompard et al., 2010). Beads were subsequently washed in wash buffer (20mM Tris pH 7.5, 50mM NaCl, 5mM MgCl2) and incubated with 1μg of actin-biotin (Cytoskeleton) / 10μl of beads for 2h at 4°C; washed in wash buffer, resuspended in Laemmli buffer and analysed by Western blotting.

For nuclear transfer experiments, sperm heads were added to egg extract supplemented as indicated, and at time points indicated, nuclei were diluted 10x in CPB buffer (50mM KCl; 20mM HEPES pH 7.6; 2% Sucrose; 5mM MgCl2) with protease inhibitors, layered onto 1ml sucrose cushion (0.7M Sucrose in CPB) and centrifuged for 5 min. at 6,000g at 4°C, and resuspended in the recipient extract. For nuclear fractionation, the pellet was further resuspended in CPB containing 0.3% Triton-X 100, then recentrifuged, supernatant recovered as nucleoplasmic fraction and pellet resuspended directly in Laemmli buffer as insoluble nuclear fraction. For immunoblot analysis, fractions corresponding to the same number of nuclei were loaded on gel.

### Immunofluorescence microscopy

Immunofluorescence microscopy using *Xenopus* egg extract nuclei and preparation of samples for visualizing actin was performed as described (Krauss *et al*, 2003). Where indicated, 20 μM biotin-dUTP or digoxigenin-dUTP (Roche), and inhibitors or DMSO, were used. Actin-Alexa Fluor and actin-biotin conjugates were obtained from Life Technologies and Cytoskeleton, respectively, and used at 25 μg/ml. DHCC was used at 2 μM. pGEX 4T1 GST-GFP-NLS plasmid was a gift from Dale Shumaker (Northwestern University, Chicago) (Moore M. S., 2000). DNA was stained with 1 μg/ml Hoechst 33258. TRITC- or rhodamine-conjugated phalloidin (Invitrogen) was used at 1/500. Secondary antibodies and Streptavidin were Alexa Fluor conjugates and were used at 1/500. Images were taken with upright Zeiss AxioimagerZ1 (100x; 1.4NA) microscope operated with Metamorph 6.2.6. software (Molecular Devices), using constant exposure time for each filter setting. Superresolution images were taken using 3D-SIM with a Deltavision OMX microscope, with Olympus UPSLAPO oil objective (100x; 1.4NA), and analyzed using OMERO.insight application. Confocal images were taken using Leica SP5-SMD microscope. The Duolink in situ PLA was performed according to the manufacturer’s instructions (Olink Bioscience, Uppsala, Sweden).

Cultured cells were seeded on gelatin-coated coverslips, synchronised and treated as described for each experiment. EdU (5-ethynyl-2´-deoxyuridine; 10 μM) or EU (5-ethynyl-uridine; 1mM for 1hr) were detected with click reaction using the Alexa Fluor^®^ 647 Imaging Kit, according to the manufacturer’s instructions (Invitrogen), and images were acquired as described above using constant exposure time between the tested conditions. For nuclear actin imaging, cells were transfected and fixed 24-48 hrs later either with 3.7% formaldehyde in cytoskeleton buffer (10 mM MES, 150 mM NaCl, 5 mM EGTA, 5 mM glucose and 5 mM MgCl2) at pH 6.2 (Small *et al*, 1999), or with glutaraldehyde essentially as described (Baarlink *et al*, 2013). TRITC-conjugated phalloidin was used at 1/1,000 for 1.5 hr. For “phalloidin alone” staining, cells were fixed with glutaraldehyde as above, and phalloidin was used at 1/200 for 20 min (for) after 3 quenching steps with sodium borohydride (1 mg/ml) (Small *et al*, 1999). Coverslips were mounted with DAPI-containing Prolong Gold or Diamond (Thermo Fisher). Image analysis, γH2A.X foci counting and signal intensity measurement was performed in Fiji-ImageJ (Schindelin *et al*, 2012) using identical parameters for all conditions. The NucleusJ plug-in (Poulet *et al*, 2015) was used to measure parameters of nuclear morphology.

For the analysis of NFκB translocation, images were acquired using a Carl Zeiss AxioimagerZ2 microscope, a plan-apochromat 40x 1.4 NA oil immersion lens and FS49 (Hoechst) and FS45 HQ (Texas Red) fluorescence filter sets, and a grid projection illumination system (aka. Apotome). The high signal to noise ratio and out of focus removal proved to be important for the analysis.

To increase the sample size, a large-field Hamamatsu Orca Flash4.0 LT sCMOS camera was used and 5x5 mosaic acquisitions were performed.

Individual tiles were analysed using a custom-designed Cell Profiler analysis routine. Briefly, nuclei masks were identified using an intensity-based automatic Otsu threshold on the Hoechst images. Cut objects at the edges of the image, as well as non-nuclear small objects were discarded. Rare, fused nuclei were segmented using an intensity algorithm. Subsequently, the nuclear masks were expanded by 10 pixels. NFKB staining integrated intensity and masked areas were then measured in both nucleus and expanded nucleus masks. Cytoplasm integrated intensities and areas were derived using expanded nucleus mask minus nucleus mask values. Mean intensity values (integrated intensity/area) and nucleus/cytoplasm mean intensity ratios were calculated.

### Statistics

Graphs were created and statistical analyses (two-tailed unpaired *t*-test) were performed in Microsoft Excel 2011 or GraphPad Prism 6. The number of cells counted in each condition and *P*-values (*, p≤0.05; **, p≤0.001; ***, p≤0.0001) are indicated in the figures. Duration of nuclear actin network and replication foci were measured manually and outliers were removed with the ROUT method (Q= 1%) in Prism 6.

### Timelapse microscopy

For live videomicroscopy, cells were seeded in glass bottom 35 mm dishes with 1 or 4 compartments, transfected as above, and image analysis was initiated 10-15 min after addition of drugs. Z-stacks (10 μm in 5 planes) were acquired every 10 minutes using an inverted microscope (Nikon) equipped with confocal spinning disk CSU-X1 Andor, 60x/1.4 oil objective using the software Andor iQ3. Stacks were processed and movies generated in Fiji. To measure the duration of nuclear actin network and PCNA foci, timelapse videos with images taken at 10-min intervals were used.

### Electron microscopy

Ten or twenty microliters of interphase *Xenopus* egg extract was supplemented with sperm DNA as described above; nuclei were allowed to assemble in the presence or absence of actin inhibitors (SMIFH2 or cytochalasin D). Sample preparation for scanning electron microscopy (SEM) was performed as described (Allen *et al*, 2007), with minor modifications. Briefly, reactions were stopped by diluting 25-fold with cold CPB buffer supplemented with protease inhibitor cocktail (Sigma-Aldrich) and centrifuged at 1,000 x g for 2 min at 4°C. Nuclei were resuspended in 0.5 ml CPB, layered onto 0.5-1 ml sucrose cushion (0.7 M in CPB) and centrifuged at 3,000 x g for 15 min at 4°C onto acetone-washed silicon chips (Agar Scientific). Nuclei were fixed in fixation buffer (80 mM PIPES, pH6.8, 30mM KCl, 1mM MgCl2, 0.25% glutaraldehyde, 2% formaldehyde, 5% w/v Sucrose) for 30 min at room temperature, washed in 0.2 M sodium cacodylate, and post-fixed with 1% osmium tetroxide solution in 0.2 M sodium cacodylate. After a wash in H2O, samples were dehydrated with increasing concentrations of ethanol (30%, 50%, 70%, 90%, and three times in absolute ethanol) followed by 10-min incubation in graded ethanol – hexamethyldisilazane. After one wash with hexamethyldisilazane, the samples were sputter-coated with approximately 3-10nm thick gold film and examined under a scanning electron microscope (Hitachi S4000 or S4800). Images were obtained using a lens detector with an acceleration voltage of 20kV at calibrated magnifications, with Axone software (version 2013; Newtec) and processed in ImageJ or Photoshop.

### Fluorescence-activated cell sorting (FACS) analysis

Cells (0.5-1 x 10^6^) were suspended in cold PBS, then pure ethanol was added to reach 70% (v/v) and fixed cells were stored in −20°C until FACS analysis. DNA was stained in PBS solution containing 2.5 μg/ml propidium iodide (PI) and 500μg/mL RNAse (Sigma-Aldrich). FACS data were obtained using FacsCalibur BD flow cytometer and visualized using Flowing software (http://www.flowingsoftware.com/ - versions 2.4.1 and above).

### Subcellular fractionation and immunoblotting

Chromatin and nucleoplasmic fractions were prepared from cell pellets essentially as described and protein concentrations were determined with the BCA method (Pierce). For immunoblotting, 10 μg of chromatin and 15 μg of soluble nuclear material were loaded on 10% or 12% polyacrylamide gels and transferred onto PVDF membranes. After blocking with 2% BSA, the corresponding antibodies were incubated for 14-16 h at 4°C.

### Mass spectrometry

Protein samples containing the nucleoskeleton and chromatin were resuspended in 2x Laemmli buffer and sonicated. Proteins (corresponding to 0.5 ml of extract) were reduced, alkylated and separated by SDS-PAGE in 4-20% gradient gels (Bio-Rad), each lane was sliced in 15 pieces and in-gel trypsin (Gold, Promega) digestion, and peptide extraction were performed essentially as described (Shevchenko *et al*, 2006). Obtained peptides were analyzed online by nano-flow HPLC-nanoelectrospray ionization using a LTQ Orbitrap XL mass spectrometer (Thermo Fisher Scientific) coupled to an Ultimate 3000 HPLC (Dionex, Thermo Fisher Scientific). Desalting and pre-concentration of samples were performed online on a Pepmap^®^ pre-column (0.3 mm x 10 mm, Dionex). A gradient consisting of 0-40% B in A for 60 min, followed by 80% B/20% A for 15 min (A = 0.1% formic acid, 2% acetonitrile in water; B = 0.1 % formic acid in acetonitrile) at 300 nL/min was used to elute peptides from the capillary reverse-phase column (0.075 mm x 150 mm, Pepmap^®^, Dionex). Eluted peptides were electrosprayed online at a voltage of 2.2 kV. A cycle of one full-scan mass spectrum (400 – 2,000 m/z) at a resolution of 60,000 (at 400 m/z), followed by 5 data-dependent MS/MS spectra was repeated continuously throughout the nanoLC separation. All MS/MS spectra were recorded using normalised collision energy (35 %, activation Q 0.25 and activation time 30 ms) with an isolation window of 3 m/z. Raw data analysis was performed using the MaxQuant software (v. 1.3.0.5). Peak lists were searched against the NCBI *Xenopus laevis* (release 130117; http://www.ncbi.nlm.nih.gov), 255 frequently observed contaminants as well as reversed sequences of all entries. The *X. laevis* genome is not fully sequenced and this results in several ‘uncharacterized proteins’. Therefore, we also searched against the *X. tropicalis* database, which is fully sequenced. The following settings were applied: spectra were searched with a mass tolerance of 7 ppm (MS) and 0.5 m/z (MS/MS). Enzyme specificity was set to Trypsin/P. Up to two missed cleavages were allowed and only peptides with at least six amino acids in length were considered. Carbamidomethylation of Cys was selected as fixed modification. Oxidation on methionine, phosphorylation on serine, threonine or tyrosine and acetylation on Protein N-term was set as a variable modification. Peptide identifications were accepted based on their false discovery rate (< 1%). Accepted peptide sequences were subsequently assembled by MaxQuant into proteins to achieve a false discovery rate of 1% at the protein level. Only proteins identified by at least 1 unique peptide or 2 peptides of at least 6 amino acids and in at least 2 of the 3 replicates were selected for further analyses.

For quantitation, the log10 of median intensity-based absolute quantification (iBAQ) was used. To compensate for the incomplete *X. laevis* database, MaxQuant .txt files were modified as follows: matchgroups were created with protein groups identified by similar peptides; GO categories for identified *X. laevis* proteins were downloaded from Uniprot after converting the GI IDs into UniprotKB accession numbers and added manually in the MaxQuant files before being loaded onto Perseus; information on “uncharacterized proteins” was extracted by assigning the UniRef90 or Uniref50 cluster. In Perseus, two groups were defined: “IBAQ D” for DMSO (control), “IBAQ P” for purvalanol A-treated (CDK-inhibited) sample. “NaN values” were converted to “0”, so that proteins identified in only one of the 2 conditions would be included in the plots. Statistical analysis was performed in Perseus selecting the 2-sample t-test with Benjamini-Hochberg and FDR 1% as parameters.

For Gene Ontology analysis in DAVID (Huang da *et al*, 2009), gene names were loaded using *Xenopus laevis* as background. All terms are presented in EV Tables but only GO BP with p-value <0.01 were further analyzed in REVIGO to remove redundant GO terms (for Figure 1B) with the following settings: SimRel method, whole Uniprot database (default settings) and Small Similarity (0.5).

## Acknowledgements

This work was supported by ANR grant ANR-09-BLAN-0252 and Ligue Nationale Contre le Cancer, ‘‘Equipe labellisée,’’ grants EL2010. LNCC/DF and EL2013. LNCC/DF and the Région Languedoc Roussillon. Thanks to the labs of J. Blow, M. Méchali, D. Görlich, B. Huelsmann, D. Maiorano and M. Hahne for providing antibodies, R. Grosse for the Lifeact-GFP-NLS, the actin-NLS R62D and all formin vectors. Thanks to B.C. Lee and V. Gladyshev for MICAL reagents. H. Hochegger, I. Hagan, P. Jay, V. Dulic, J. Hutchins and A. Crevenna for comments on the manuscript. Thanks to M. Goldberg for advice on sample preparation for electron microscopy, C. Cazevieille of the CRIC-IURS, D. Cot of the IEM, V. Georget, J. M. Langerak and J. Cau of MRI imaging facilities, Montpellier, for assistance with electron and light microscopy and analysis. Thanks to R. Audo for help with NF-κB experiments.

## Author Contributions

NP and LK conceived, performed and analysed most experiments and wrote the manuscript. BH performed other experiments. SU and NP performed proteomics. MR co-supervised NP for proteomics. AC provided an important intellectual contribution. JD supervised NPE experiments. NM co-supervised nuclear transport experiments. DF conceived and directed the study and wrote the manuscript.

## Conflict of Interest Statement

The authors declare no competing interests.

## Supplementary information

Supplementary information includes 6 videos and 4 tables.

## Supplementary Figure legends

**Supplementary Figure 1.**
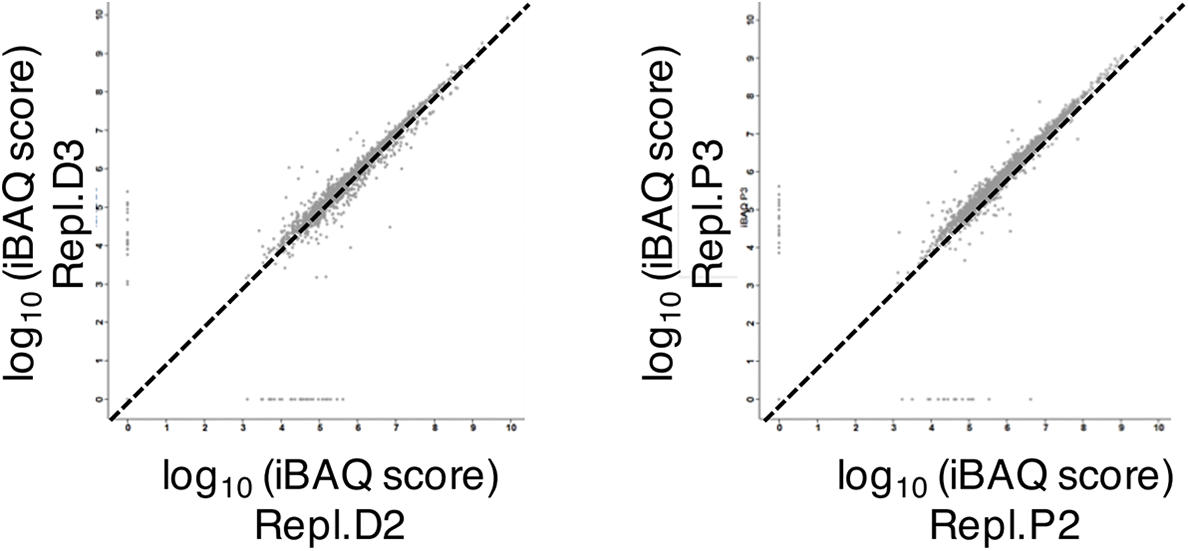
Scatter plots showing the high reproducibility of MS/MS runs between replicates.

**Supplementary Figure 2.**
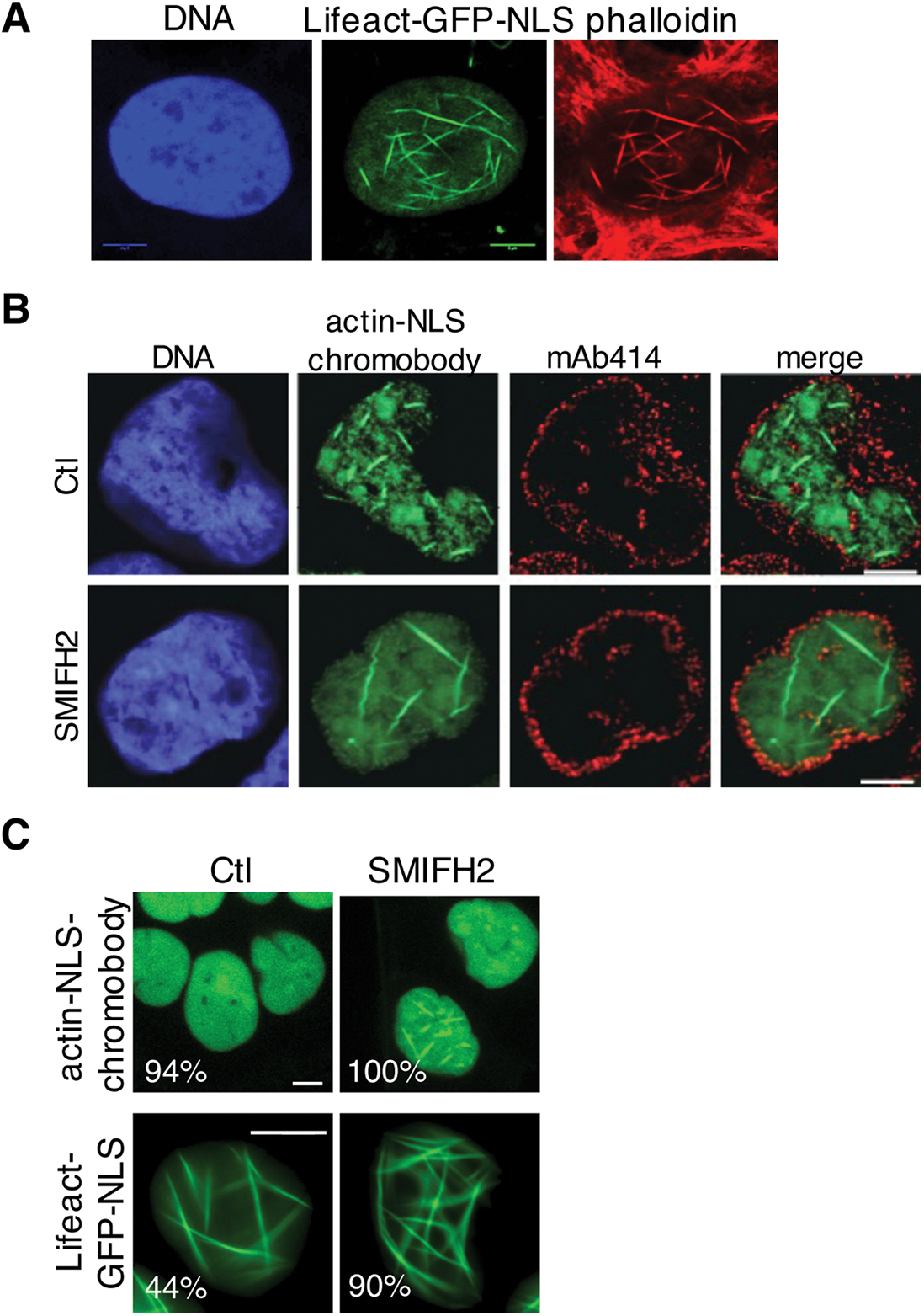
Effect of actin drugs and probes on nuclear actin dynamics and form in human cells. **A** U2OS cells transiently expressing Lifeact-NLS-GFP, fixed and stained with phalloidin and DAPI (DNA). Bar, 5 μm. **B** Interphase U2OS cells expressing actin-NLS chromobody co-stained with mAb414 antibody and DAPI treated with DMSO (Ctl) or SMIFH2 (50 μΜ) for 2 hours. Bar, 5μm. **C** Snapshots of live U2OS cells expressing actin-NLS chromobody or Lifeact-GFP-NLS in the presence of DMSO (Ctl) or SMIFH2 (50μΜ). Numbers represent the percent of cells showing the presented phenotype; n>100 cells in each condition from ≥3 independent experiments. Bar, 5 μm.

**Supplementary Figure 3.**
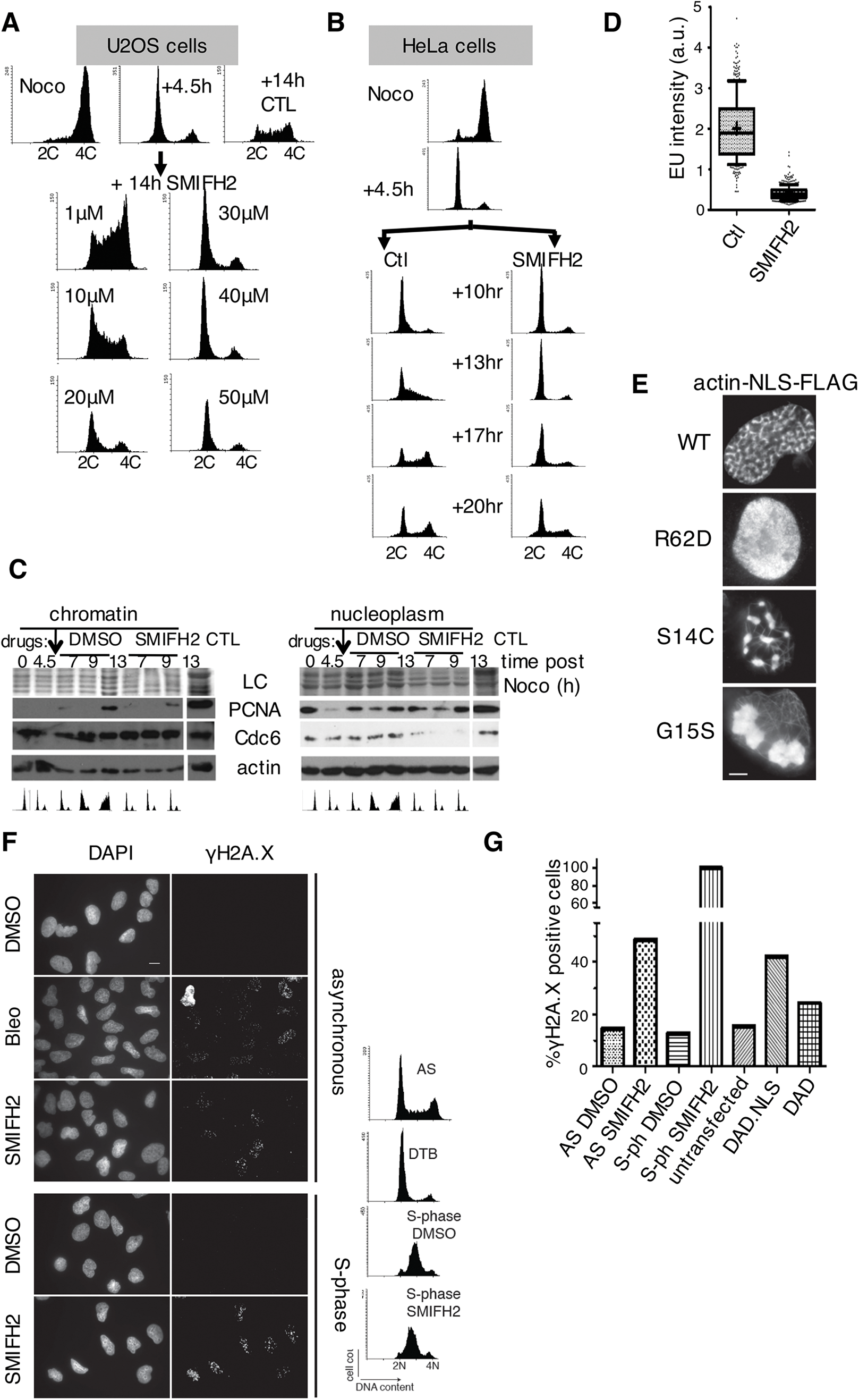
Disrupting actin dynamics hinders DNA replication and causes replication stress in human cells. **A** FACS analysis of G1-synchronised U2OS cells in the presence of increasing concentrations of SMIFH2. Cells were collected when control cells were in S-phase (+14h). **B** FACS analysis of HeLa cells, synchronised in G1 as in Fig 4A, and treated with DMSO (Ctl) or SMIFH2, collected at the time points indicated. **C** Immunoblotting of chromatin and nucleoplasmic fractions from cells in Fig 4A. FACS profiles for each time-point are shown. Ctl, U2OS cell lysate; LC, loading control. **D** Quantification of EU signal intensity from experiment presented in Fig. 2b (n>400). **E** Immunofluorescence images of U2OS cells transfected with the indicated FLAG-tagged actin-NLS constructs. DNA was stained with DAPI, actin-NLS constructs with anti-Flag antibody. Bar, 5μm. **F** Asynchronous (AS) or double-thymidine block (DTB) S-phase-synchronised U2OS cells were treated for 2hrs with DMSO, bleomycin (Bleo), or SMIFH2, and stained for γH2A.X. Corresponding FACS profiles are shown. **G** Quantification of γH2A.X-positive cell number from experiment presented in F, and in cells transiently transfected with mDia2 DAD.NLS or DAD constructs. Cells with >10 foci were considered positive.

**Supplementary Figure 4.**
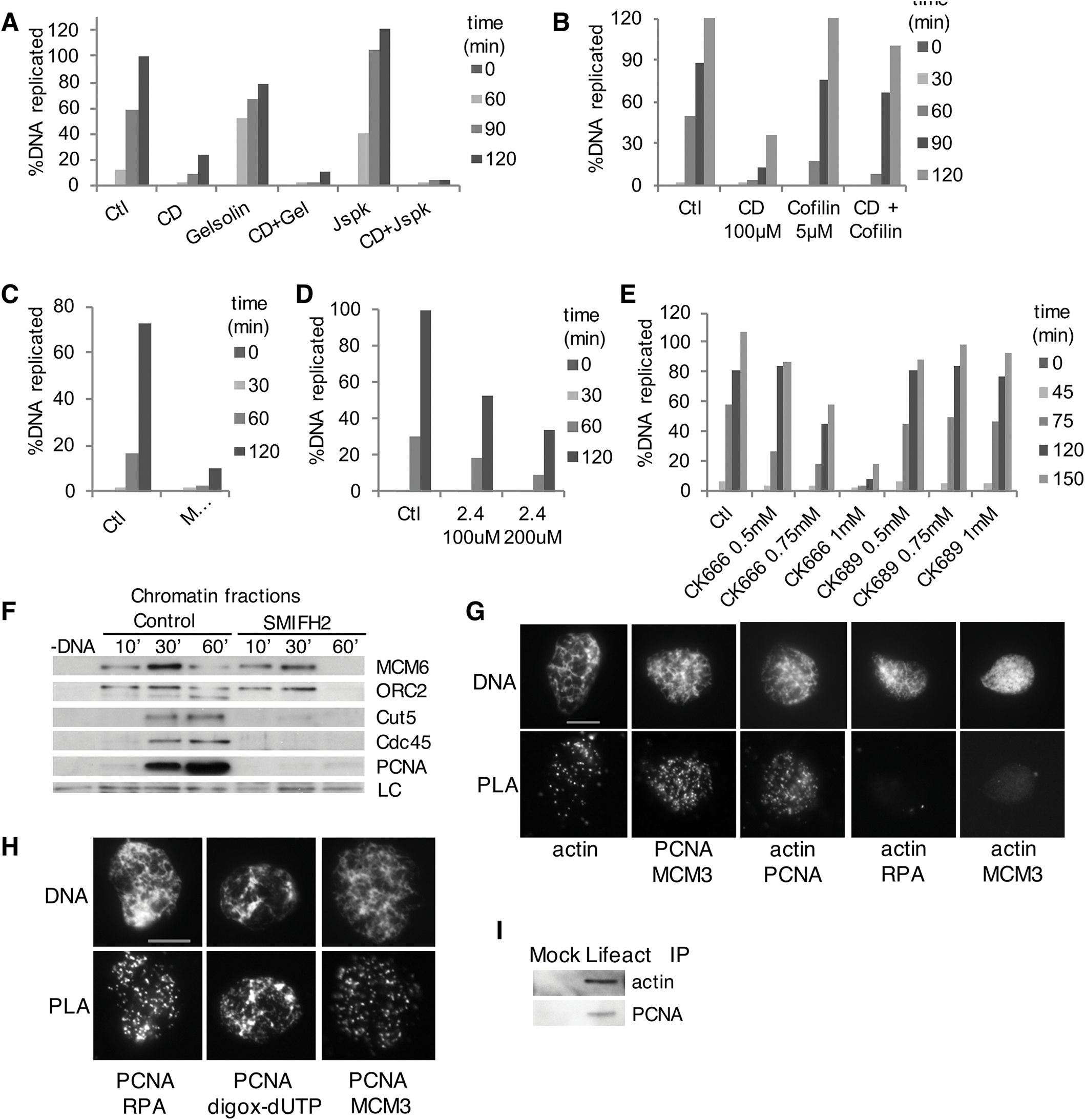
Effect of actin drugs and actin regulators on DNA replication in *Xenopus* egg extracts. **A** Replication time-course of sperm chromatin in extract treated with Cyt D (CD), gelsolin, Cyt D and gelsolin (CD+Gel), jasplakinolide (Jspk), or Cyt D and jasplakinolide (CD+Jspk). **B-E** Replication assays in control extract or in extracts supplemented with Cyt D (CD) with or without cofilin (**B**); recombinant MICAL2 protein (**C**), formin inhibitor 2.4 (**D**), or Arp2/3 inhibitor CK666 or its inactive analogue CK689 (**E**). **F** Chromatin loading of pre-RC and pre-IC factors in control conditions or in the presence of SMIFH2 (500μM). LC, loading control. **G, H** Interaction of endogenous proteins indicated and biotin-dUTP was probed by PLA in nuclei formed in control XEE for 60 min. Primary anti-actin rabbit antibodies were combined with secondary rabbit PLA probes. Bar, 10μm. **I** Nuclei formed in control extract, were lysed at 60 min and actin was precipitated using biotinylated Lifeact peptide immobilised on streptavidin beads (Mock IP, no peptide); beads were blotted for actin and PCNA.

**Supplementary Figure 5.**
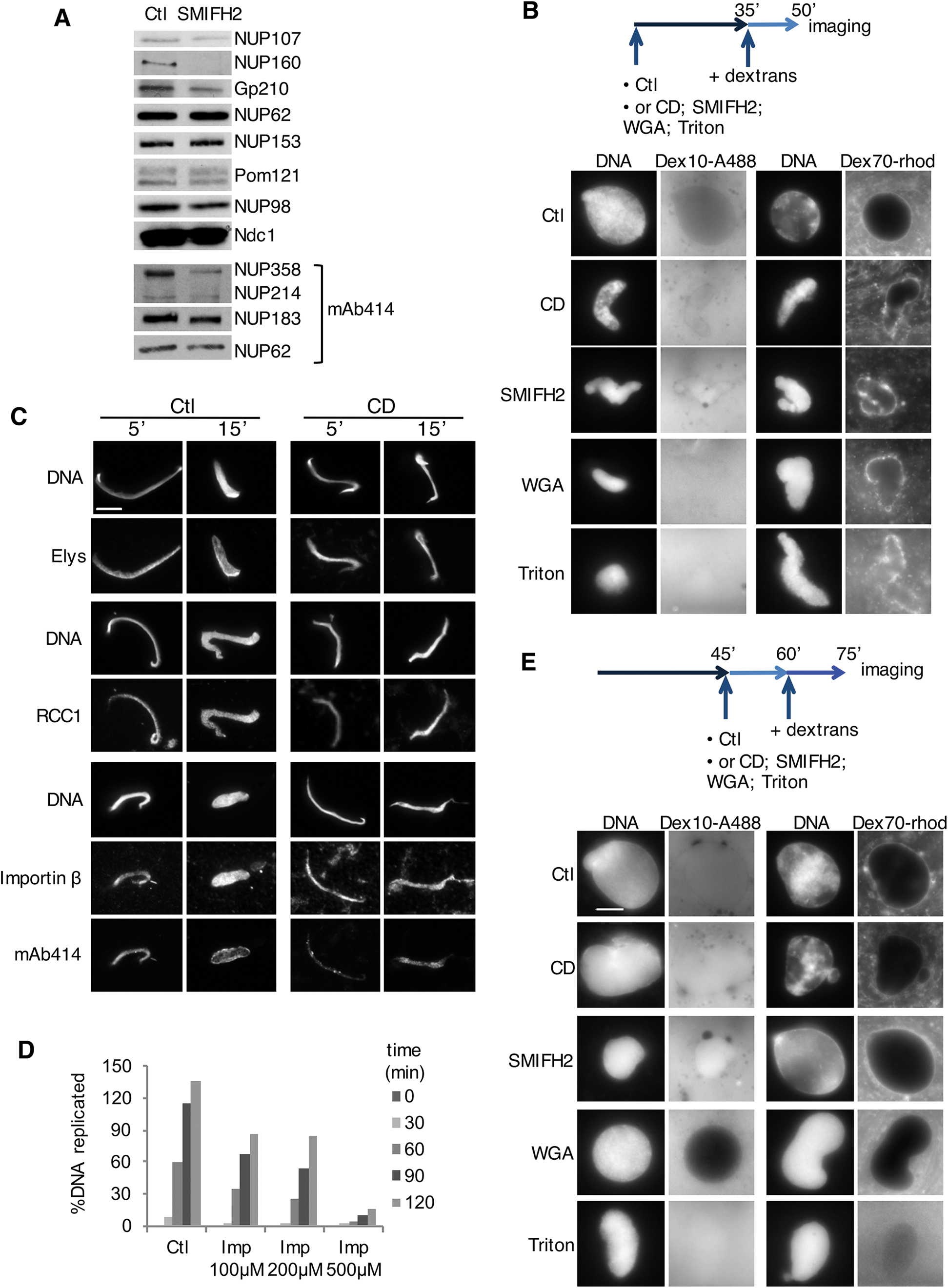
Active nuclear transport requires actin dynamics. **A** Western blots of total nuclear fractions in control and formin-inhibited (SMIFH2) extracts, probed with antibodies to the NUPs indicated; the same number of nuclei was analysed for each condition. **B** Scheme: Nuclei were formed in control extract or in the presence of Cyt D (CD), SMIFH2, WGA or Triton; at 35 min. Dextran10-Alexa488 (Dex10-A488) or Dextran70-rhodamine (Dex70-rhod) were added; nuclei were imaged directly at 50 min. Bar, 10 μm. **C** Immunofluorescence images of nuclei formed for 5 and 15 min in control or cytochalasin D-treated extracts, stained for DNA, Elys, RCC1, importin-β and FG-NUPs (mAb414). Bar, 10 μm. **D** Replication time course of sperm chromatin in extract treated with importazole (Imp) at the concentrations indicated. **E** Scheme: Nuclei were formed in control extract; at 45 min Cyt D (CD), SMIFH2, WGA or Triton were added, followed by addition of Dextran10-Alexa488 (Dex10-A488) or Dextran70-rhodamine (Dex70-rhod); nuclei were imaged directly at 75 min. Bar, 10 μm.

**Supplementary Figure 6.**
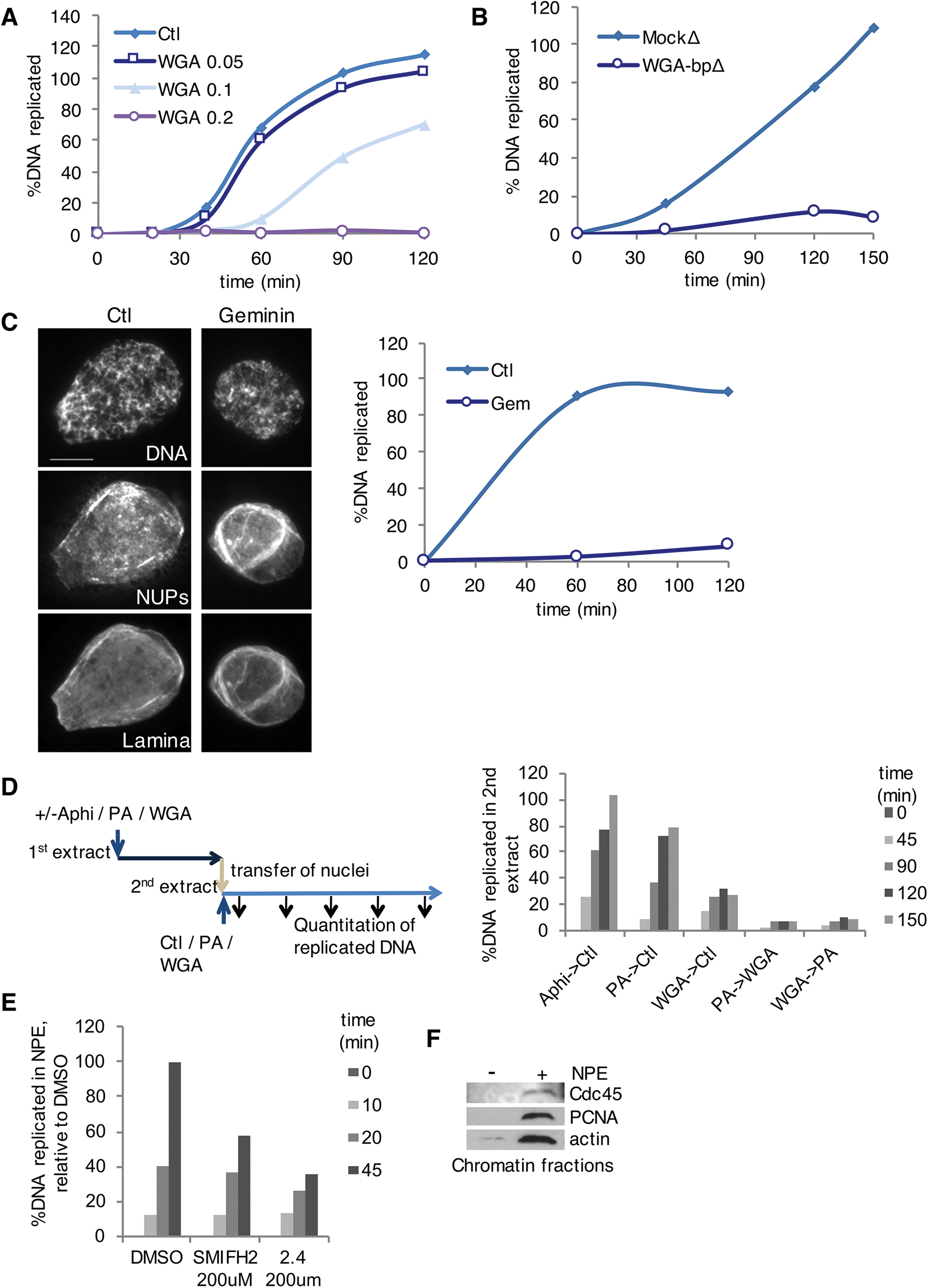
Ongoing nuclear transport is required to promote DNA replication. **A** Replication assay in control extract and in extract supplemented with WGA at the concentrations indicated (mg/ml). **B** Replication assay of Mock- and WGA-binding protein-depleted (WGA-bp∆) extract. **C** Left, immunofluorescence images of nuclei formed for 60 min in control and geminin(40nM)-containing extracts, stained for DNA, NUPs (mAb414) and Lamin B3. Bar, 10μm. Right, replication time-course of control and geminin-treated extract. **D** Scheme: reciprocal nuclear transfer experiment, in which first extracts contained aphidicolin (Aphi) and either PA or WGA, or were control; nuclei were isolated after 45 min and transferred to the alternative condition; DNA replication was assayed in the second extract. **E** Replication time course of sperm chromatin in NPE extract treated with SMIFH2 or 2.4 formin inhibitor (mean values of two replicates), both used at 200μM. **F** Western blot for indicated proteins of chromatin fractions with or without addition of NPE.

Table S1: Entire MS/MS Dataset. Proteins identified in the nuclear structural proteome of control and CDK-inhibited extracts.

Table S2: GO Annotations of the nuclear structural proteome using DAVID.

**Table S3.**
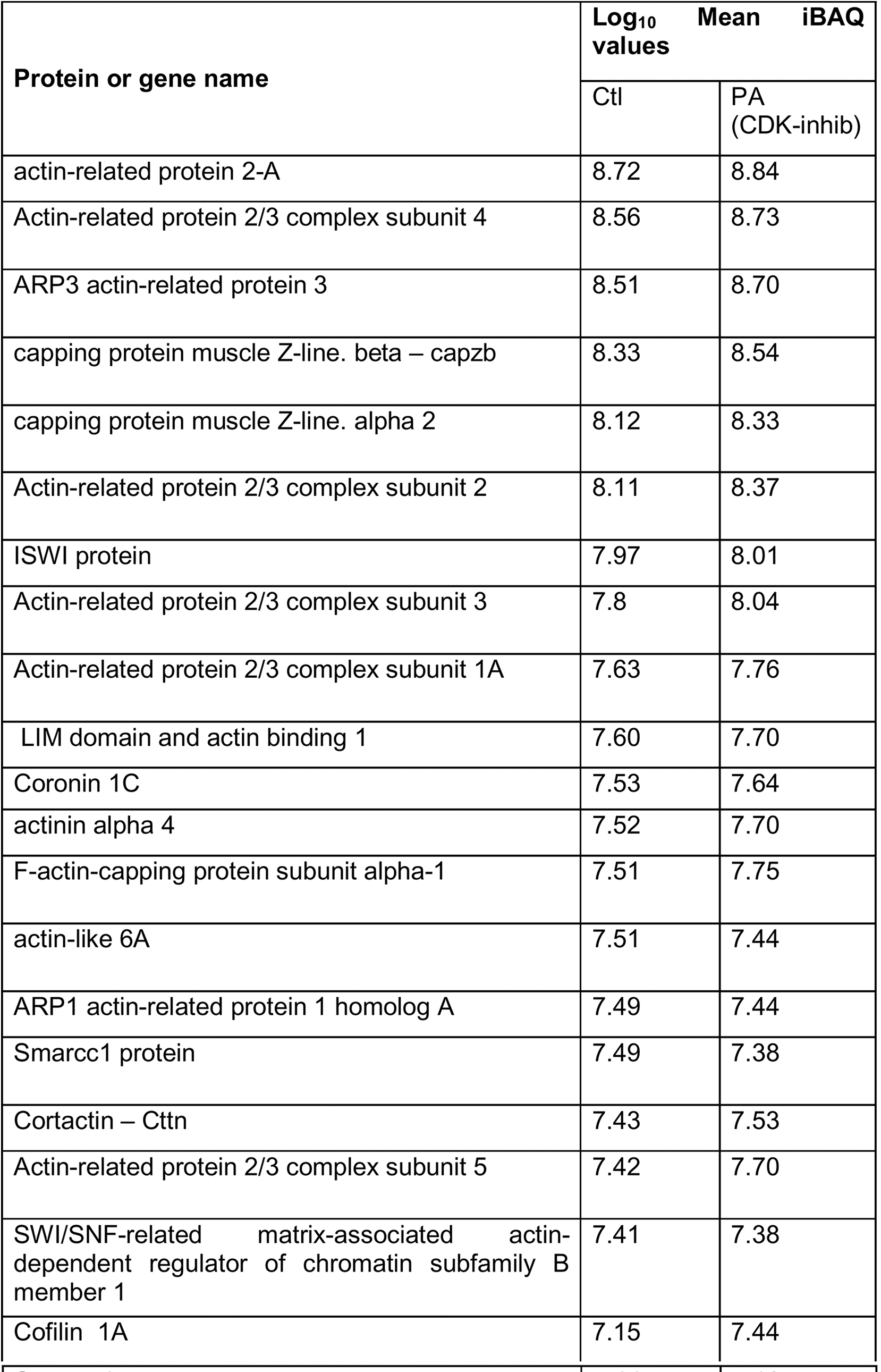
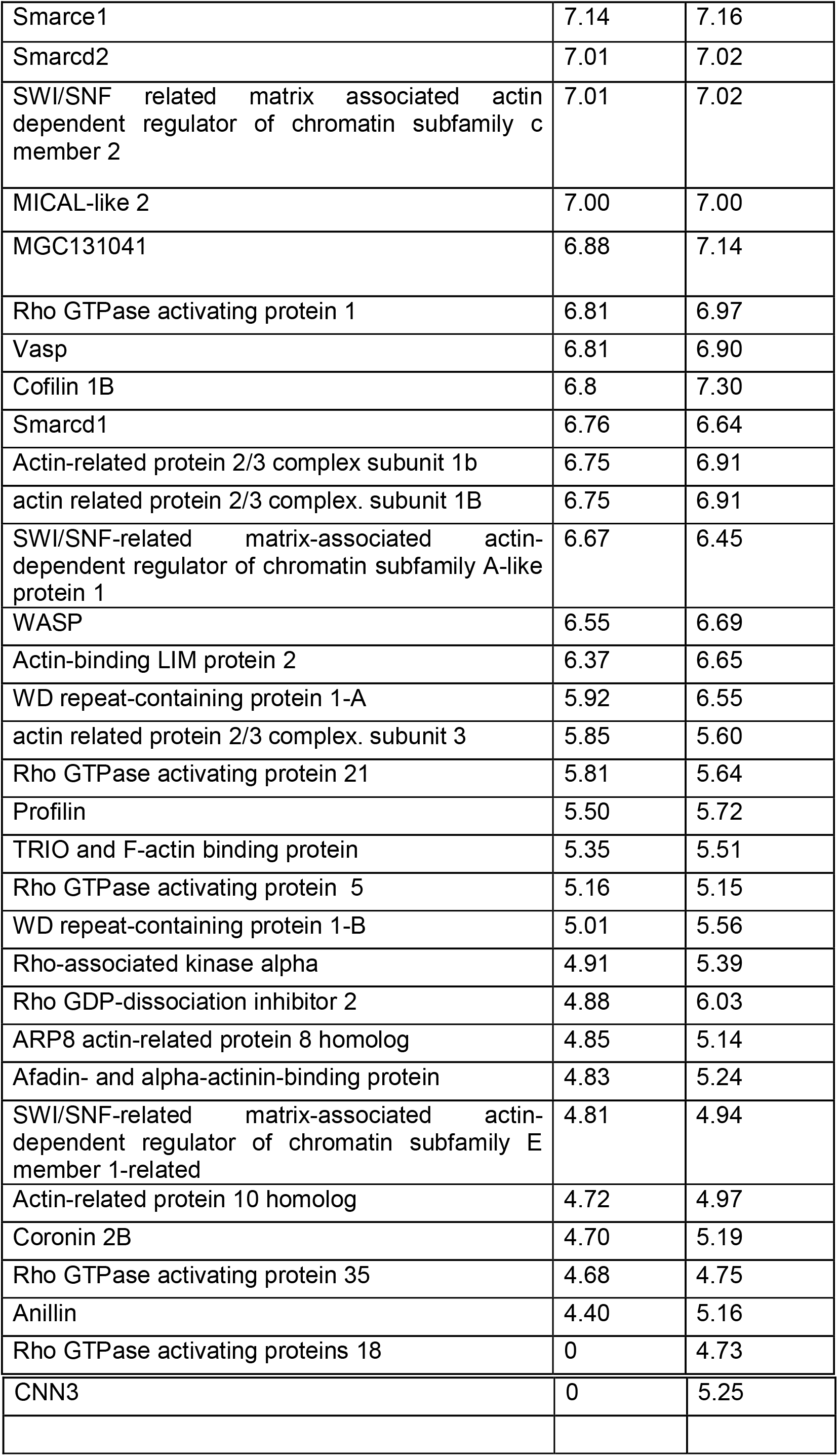
Actin regulators identified in the structure proteome of nuclei.

**Table S4:** List of formins in the structure proteome of nuclei in our dataset (*X. tropicalis* database)

| Gene names | Fasta headers | No of peptides |  |
| --- | --- | --- | --- |
|  |  | Ctl | CDK-inh |
| diaph1 | F6UXX7_XENTR Uncharacterized protein OS=Xenopus tropicalis | 4 | 4 |
| diaph3 | F6VAH9_XENTR Uncharacterized protein (Fragment) OS=Xenopus tropicalis | 3 | 3 |
| fmn2 | F6UL32_XENTR Uncharacterized protein OS=Xenopus tropicalis | 2 | 3 |

**Movie 1 Visualization of early G1 nuclear actin filaments in live human cancer cells.**

Timelapse confocal microscopy of U2OS cells stably expressing actin-NLS chromobody (green) and transiently transfected with PCNA chromobody (red).Numbers denote time in hr:min; bar, 5 μm.

**Movie 2 Prolonged expression of Lifeact-GFP-NLS stabilises nuclear actin filaments and impairs cell cycle progression.**

Timelapse confocal microscopy of U2OS cells transiently expressing Lifeact-GFP-NLS. Numbers denote time in hr:min; bar, 5μm.

**Movie 3 Expression of mDiaDAD.LG in the nucleus impairs replication foci formation.**

Timelapse confocal microscopy of S-phase U2OS cells stably expressing the PCNA chromobody (red) and transiently transfected with the mDiaDAD.LG.NLS construct (green). Numbers denote time in hr:min; bar, 5μm.

**Movie 4 SMIFH2 impairs nuclear actin dynamics.**

Timelapse confocal microscopy of U2OS cells stably expressing actin-NLS chromobody (green), in the presence of SMIFH2 (50 μM). Numbers denote time in hr:min; bar, 5μm.

**Movie 5 SMIFH2 stabilises early G1 nuclear actin filaments.**

Timelapse confocal microscopy of early G1 U2OS cells stably expressing actin-NLS chromobody (green), in the presence of SMIFH2 (50 μM). Numbers denote time in hr:min; bar, 5 μm.

**Movie 6 Perturbing nuclear actin dynamics abolishes replication foci formation.**

Timelapse confocal microscopy of S-phase U2OS cells stably expressing actin-NLS chromobody (green) transiently transfected with the PCNA chromobody (red), in the presence of SMIFH2 (50 μM). Numbers denote time in hr:min; bar, 5 μm

